# A dual-strategy expression screen for candidate connectivity labels in the developing thalamus

**DOI:** 10.1101/079327

**Authors:** Olivia Bibollet-Bahena, Tatsuya Okafuji, Karsten Hokamp, Kevin J. Mitchell

## Abstract

The thalamus or “inner chamber” of the brain is divided into ~30 discrete nuclei, with highly specific patterns of afferent and efferent connectivity. To identify genes that may direct these patterns of connectivity, we used two strategies. First, we used a bioinformatics pipeline to survey the predicted proteomes of nematode, fruitfly, mouse and human for extracellular proteins containing any of a list of motifs found in known guidance or connectivity molecules. Second, we performed clustering analyses on the Allen Developing Mouse Brain Atlas data to identify genes encoding surface proteins expressed with temporal profiles similar to known guidance or connectivity molecules. In both cases, we then screened the resultant genes for selective expression patterns in the developing thalamus. These approaches identified 82 candidate connectivity labels in the developing thalamus. These molecules include many members of the Ephrin, Eph-receptor, cadherin, protocadherin, semaphorin, plexin, Odz/teneurin, Neto, cerebellin, calsyntenin and Netrin-G families, as well as diverse members of the immunoglobulin (Ig) and leucine-rich receptor (LRR) superfamilies, receptor tyrosine kinases and phosphatases, a variety of growth factors and receptors, and a large number of miscellaneous membrane-associated or secreted proteins not previously implicated in axonal guidance or neuronal connectivity. The diversity of their expression patterns indicates that thalamic nuclei are highly differentiated from each other, with each one displaying a unique repertoire of these molecules, consistent with a combinatorial logic to the specification of thalamic connectivity.

## Introduction

The thalamus, or “inner chamber” of the brain, is a crucial nexus in the brain’s circuitry. It is not only a relay station that conveys sensory information from the periphery to the cerebral cortex, it is also a conduit for cortico-cortical communication [1], as well as a central node in pathways controlling action selection, through cortico-baso-thalamo-cortical loops [2]. In addition, the thalamus is interconnected with many other brain structures, including hippocampus, hypothalamus, amygdala, inferior and superior colliculus [3], cerebellar nuclei [4], substantia nigra, brainstem, spinal cord and many others. The thalamus proper, or dorsal thalamus, is also intimately interconnected with the prethalamic reticular nucleus [5], which provides inhibitory regulation of information flow through thalamus.

A striking characteristic of the thalamus is its subdivision into ~30 discrete nuclei, which subserve distinct functions and which have highly selective connectivity patterns with the structures mentioned above, most obviously with specific cortical areas. Some nuclei, such as those conveying primary sensory information, project quite selectively to one or a small number of cortical areas, while others, which integrate signals from multiple sources, project in a more diffuse manner across larger areas of cortex. Within each nucleus there are also varying proportions of distinct cell types that either project to specific areas, driving receptive field properties in input layers of cortex, such as layer 4, or that project more widely across cortex and provide modulatory inputs, for example to layer 1 [6,7].

Different thalamic nuclei thus act as discrete targets for innervation from a wide array of sources, and, in turn, project their axons to numerous other structures with areal, laminar and cell-type specificity. In order to establish these connections, growing axons must be guided to appropriate regions, must recognise the appropriate target and avoid inappropriate ones, and must make the right kinds of synapses on the right kinds of cells. These processes are mediated in general by surface and secreted proteins, which act as signals and receptors, enabling cellular recognition for pathfinding and target selection.

Numerous protein families have been identified as playing important roles in thalamic axon guidance and connectivity. Many of these, including Ephrins/Eph-receptors, Netrins/DCC/Unc5s, Slits/Robos, Neuregulin-1/Erbb4, secreted semaphorins/neuropilins and L1 cell adhesion molecules, mediate general processes such as avoidance of the hypothalamus, projection into the internal capsule and topographic organisation of thalamocortical and/or corticothalamic projections through this intermediate target region [reviewed in 8,9]. By contrast, very few molecules have been found so far that mediate more specific connectivity relationships of particular thalamic nuclei, such as Cdh6 [10], or that control sub-organisation of projections within thalamic nuclei, such as Lrp8 [11] or Ten-m3/Odz3 [12], which also regulates topography of projections to striatum [13].

At earlier stages, a number of studies have described the combinatorial expression patterns of patterning molecules or transcription factors across the thalamus, many of which are expressed selectively in some nuclei and not others [14-18]. Some have also documented the differential expression of selected surface proteins across the thalamus [15,19,20], and in some cases direct links have been shown between transcription factor and surface molecule expression [21,22]. However, these analyses have not been performed in a systematic or comprehensive fashion. Thus, while we have learned a lot about how the developing thalamus is patterned and how the fates of different nuclei are specified, we still know relatively little about the combinatorial code of surface molecules that specifies nuclear connectivity.

Here, we describe two parallel approaches to identify genes encoding surface or secreted proteins expressed in discrete patterns across the developing thalamus. First, we screened the predicted proteomes of human, mouse, fly and worm for conserved genes encoding predicted transmembrane proteins with any of a number of protein motifs commonly found in axon guidance molecules. These genes were screened by *in situ* hybridization to find those expressed in selective or differential patterns across the neonatal thalamus. Second, we analysed the Allen Developing Mouse Brain Atlas (devABA) database for genes encoding surface or secreted proteins, which showed a similar temporal expression profiles similar to known genes for axon guidance or synaptic connectivity. The expression patterns of these genes were then examined to identify selectively or differentially expressed genes at mid or late embryonic stages. Together, these approaches have identified 82 genes encoding candidate connectivity labels in the developing thalamus. The expression patterns of these genes are highly diverse, such that individual thalamic nuclei express distinct repertoires of these surface molecules, consistent with a combinatorial logic to the specification of connectivity.

## Results

### A bioinformatics and expression screen to identify conserved candidate connectivity labels

The proteomes of mammals contain many predicted proteins of unknown function. We were interested to discover genes encoding predicted transmembrane proteins that contain any of a number of protein motifs found in known axon guidance molecules. As many known axon guidance molecules are highly conserved from vertebrates to invertebrates, we further concentrated on proteins that had predicted orthologues in the fruitfly Drosophila melanogaster and/or in the nematode Caenorhabditis elegans.

To identify such proteins, we used protein localisation and motif searches to annotate the predicted proteomes of worm, fly, mouse and human. We also generated a matrix of pairwise BLAST scores across all the members of these proteomes and clustered them using the TRIBE-MCL algorithm, as previously described [23]. We screened through the resultant outputs to find clusters with mammalian and fly or worm members encoding predicted transmembrane proteins, which also contained any of the following motifs: immunoglobulin (Ig) domain, fibronectin type 3 domain, cadherin domain, leucine-rich repeat (LRR), EGF repeat, CUB domain, sema domain, or plexin repeat. We were particularly interested in discovering novel axon guidance or connectivity cues and so excluded genes or gene families where such a function had already been demonstrated in the mouse at the time the screen was performed. Using this approach, we prioritized a set of 42 genes for expression screening (Table S1).

We performed *in situ* hybridization for these genes on P0 mouse brain sections, as previously described for genes encoding LRR proteins [23]. Here we concentrate on members of other gene families that show selective or differential expression across the thalamus (Table 1 Column A).

**Table 1.**
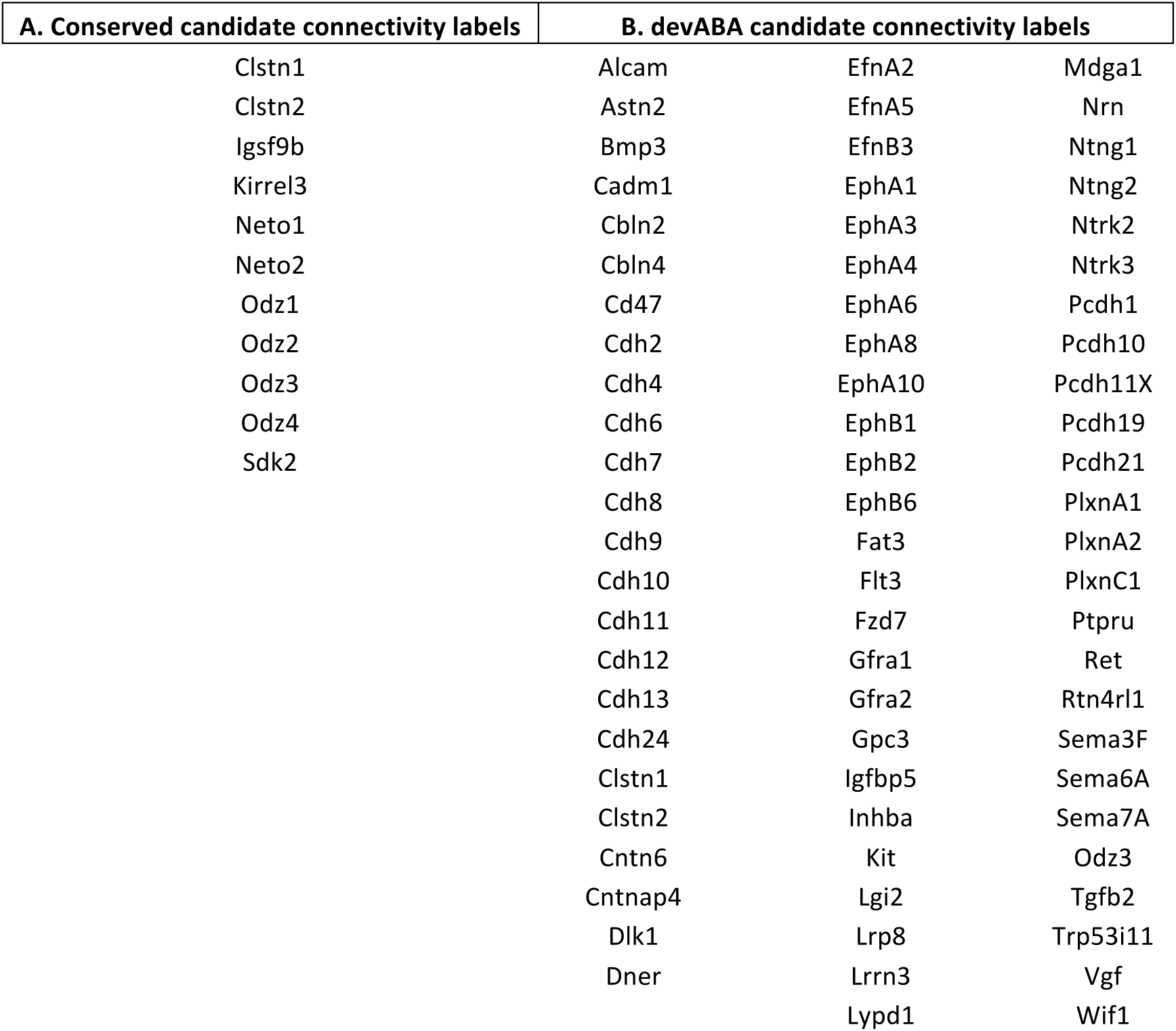
**Candidate connectivity labels**. This table lists all 82 genes for which we present expression patterns. Clstn1, Clstn2 and Odz3 appear on both lists.

***Immunoglobulin superfamily members***: We found three Ig superfamily members with selective expression across the thalamus: Igsf9b, Kirrel3, and Sdk2 (Figure 1; Figure S1). Igsf9b (Immunoglobulin Superfamily, Member 9B) is an orthologue of the Drosophila gene *turtle*, which has been implicated in various aspects of axon guidance [reviewed in 24]. Both Igsf9b and Igsf9 have recently been shown to play roles in inhibitory synapse development in mammals [25,26]. Igsf9b is more strongly expressed in some thalamic nuclei than others, but we detected only weak expression of Igsf9 in thalamus (not shown).

**Figure 1.**
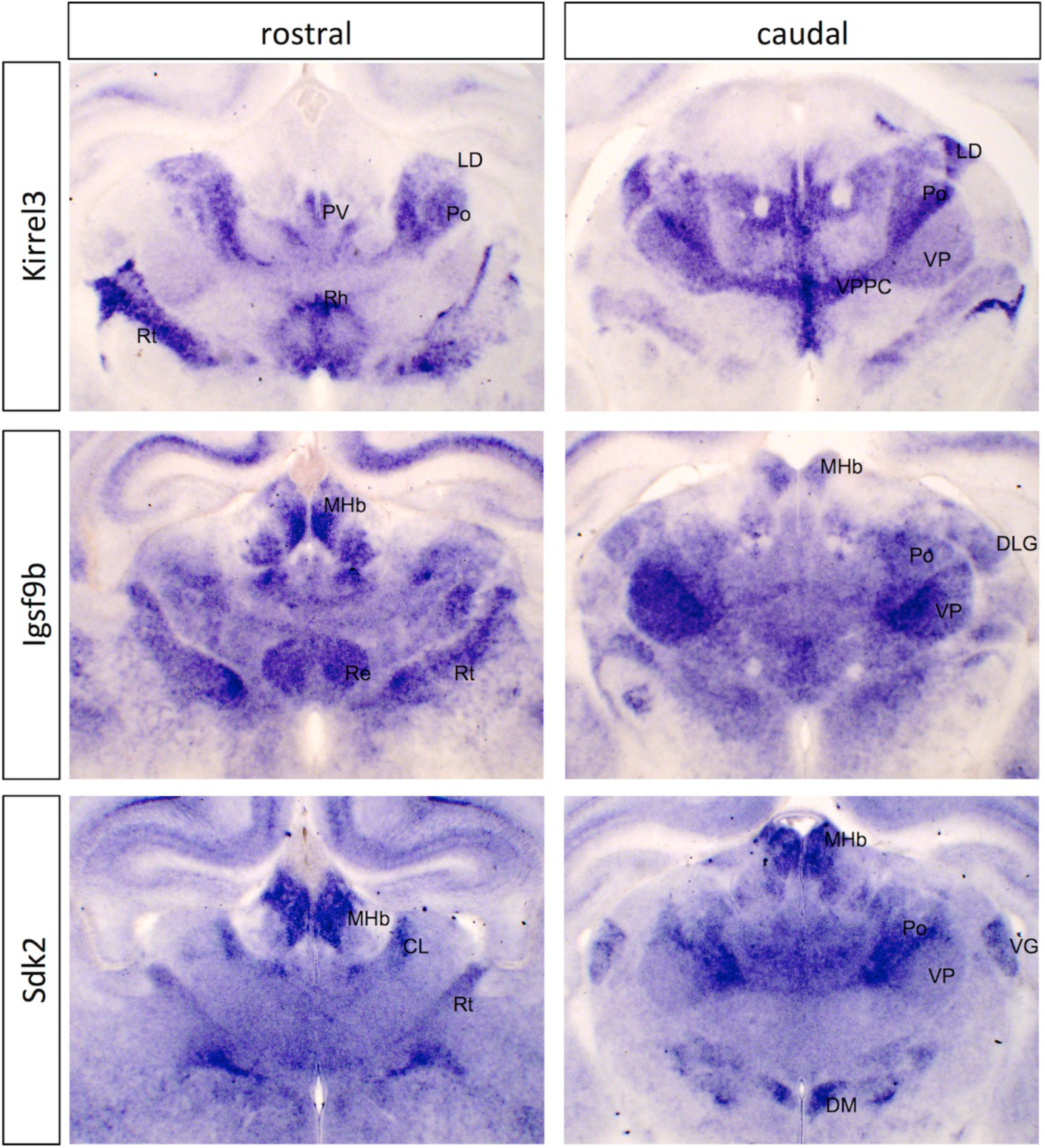
***In situ* hybridization patterns for immunoglobulin superfamily genes in dorsal thalamus at P0.** Two coronal sections are shown for Kirrel3, Igsf9b and Sdk2, one rostral and one more caudal. The entire corresponding sections are shown in Figure S1, for context. CL, centrolateral nucleus; DLG, dorsolateral geniculate nucleus; DM, dorosmedial hypothalamic nucleus; LD, laterodorsal nucleus; MHb, medial habenula; Po, posterior thalamic nuclear group; PV, paraventricular nucleus; Re, reuniens nucleus; Rh, rhomboid nucleus; Rt, reticular nucleus; VG, ventral geniculate nucleus; VP, ventral posterior nucleus; VPPC, ventral posterior nucleus parvicellular part.

Kirrel3 (Kin Of IRRE Like 3 (Drosophila), also known as NEPH2) is an orthologue of the fly gene *irregular chiasm* (*IrreC*), which functions in axonal pathfinding and target selection [reviewed in 27] and the C. elegans gene *SYG-1*, which is implicated in selective synaptogenesis [28]. Kirrel3 has recently been implicated in axon guidance in the vomeronasal system [29] and synapse formation in the hippocampus [30], but functions in the developing thalamus have not been described. Kirrel1 and Kirrel2 both showed only weak/background expression in thalamus (not shown).

Sdk2 (Sidekick2) is an orthologue of the fly gene *sidekick*, which regulates cellular differentiation in the fly eye [31]. An important role for Sidekicks has also been demonstrated in the vertebrate retina in specifying lamina formation and synaptic connectivity [32,33]. Sdk2 is quite selectively expressed across thalamic nuclei while Sdk1 is expressed at high levels across the whole dorsal thalamus (not shown).

***Odz (Teneurin) genes***: We found that all four members of the Odz/Teneurin family are expressed in selective fashion across the developing thalamus at P0 (Figure 2; Figure S2) and at E15 (Figures S3–S4). These genes are orthologues of the fly *odd oz/teneurin* genes *ten-a* and *ten-m,* which function in synaptic connectivity in the olfactory and neuromuscular systems [reviewed in 34]. In mammals, they have both homophilic and heterophilic interactions. Odz2 (Ten-m2) and Odz4 (Ten-m4) are expressed in similar, but not identical patterns, and Odz3 (Ten-m3) expression overlaps substantially with them. Odz1 (Ten-m1) is expressed in a broadly complementary pattern to the other three genes. These genes also show graded expression across cortex and striatum at E15 and P0 (Figures S2 and S4). Roles for Odz2 and Odz3 have been demonstrated in establishing the topography and segregation of ipsilateral and contralateral afferents in the dLGN and of thalamic afferents in visual cortex [reviewed in 35]. More recently, Odz3 has also been shown to control topography of thalamostriatal projections [13].

**Figure 2.**
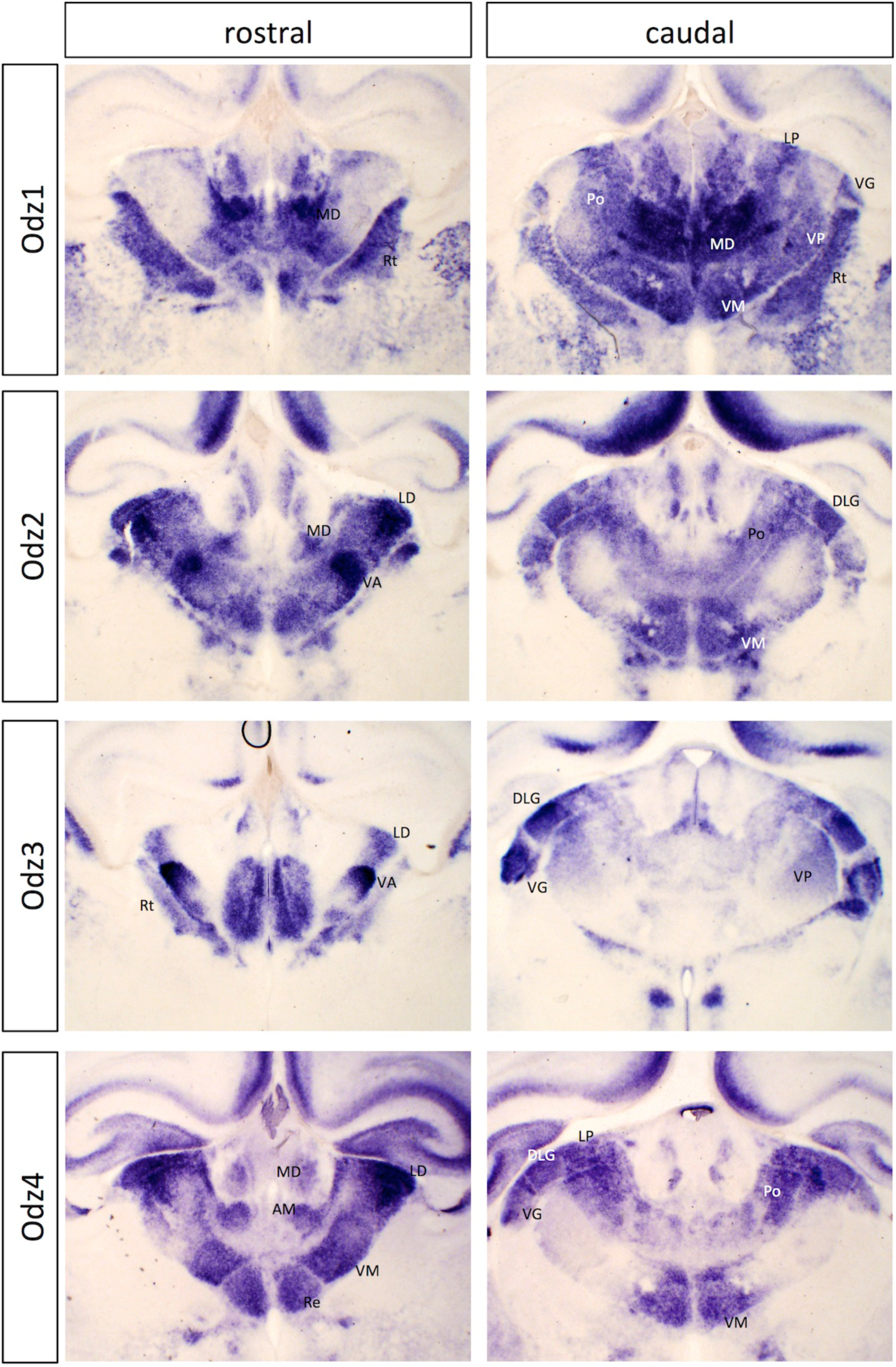
***In situ* hybridization patterns for Odz family genes in dorsal thalamus at P0.** Two coronal sections are shown for Odz1, Odz2, Odz3 and Odz4, one rostral and one more caudal. The entire corresponding sections are shown in Figure S2, for context. AM, anteromedial nucleus; DLG, dorsolateral geniculate nucleus; LD, laterodorsal nucleus; MD, mediodorsal nucleus; Po, posterior thalamic nuclear group; Re, reuniens nucleus; Rt, reticular nucleus; VA, ventral anterior nucleus; VG, ventral geniculate nucleus; VM, ventromedial nucleus; VP, ventral posterior nucleus.

***Neto genes***: The neuropilin and tolloid-like genes Neto1 and Neto2, orthologues of the fly gene *neto*, also showed discrete expression in the thalamus, with Neto2 more widely expressed (Figure 3; Figure S5). In some parts of the thalamus, their expression was strikingly complementary, particularly in the ventrobasal complex, which expresses high levels of Neto1 but not Neto2, while many other nuclei show the opposite pattern. The Neto proteins function as auxiliary subunits for kainate receptors in mammals [reviewed in 36] and in glutamate receptor clustering at the neuromuscular junction in flies [37]. No role in axon guidance or synaptic connectivity has been demonstrated for these genes, but their early expression (Figure S6) and protein similarity to neuropilins would certainly be consistent with such a function.

**Figure 3.**
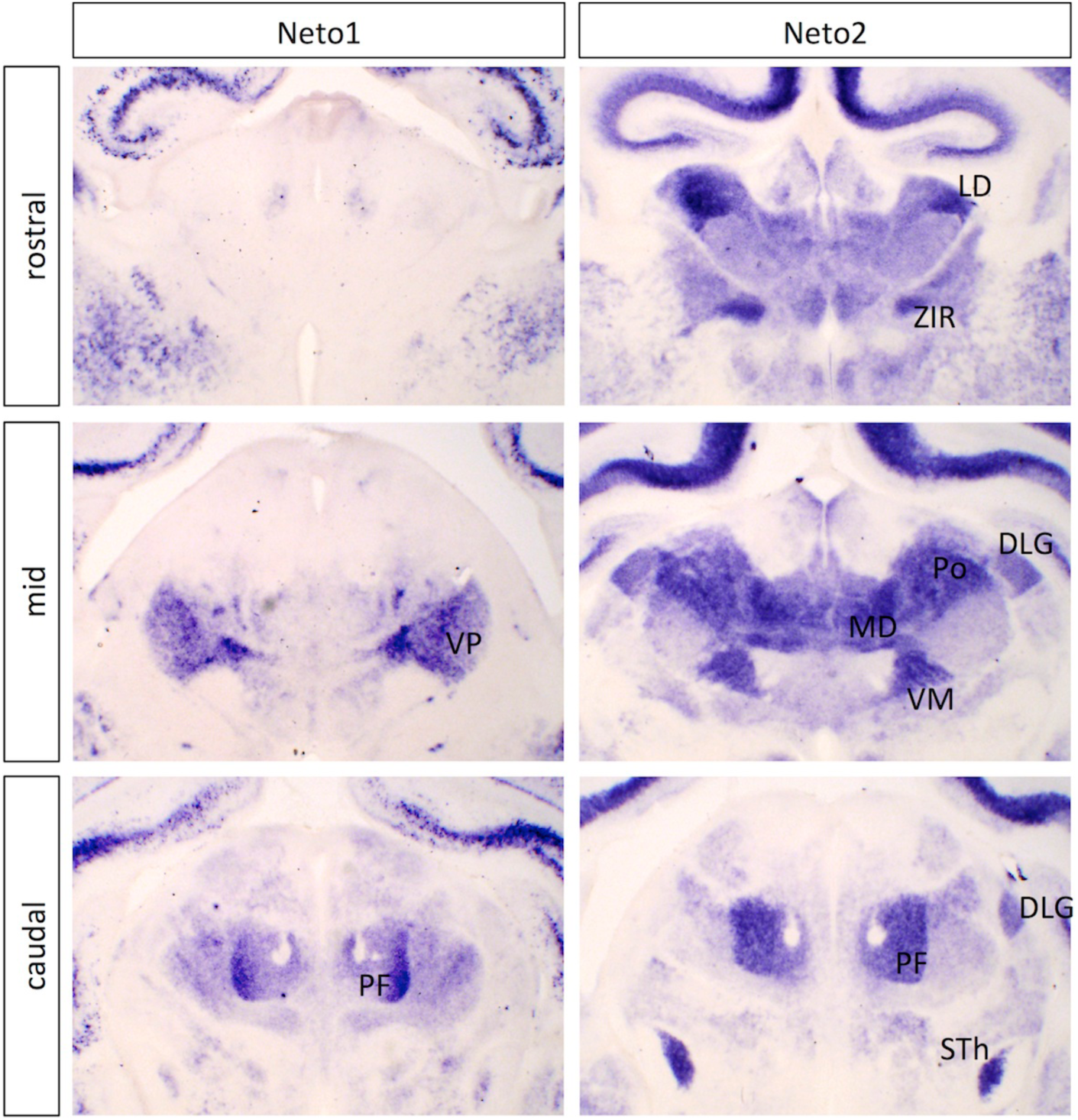
***In situ* hybridization patterns for Neto family genes in dorsal thalamus at P0.** Three coronal sections are shown for Neto1 and Neto2, one rostral, one at an intermediate level (mid) and one more caudal. The entire corresponding sections are shown in Figure S5, for context. Expression of the Neto genes is notably complementary in some places (compare mid sections) and overlapping in others (e.g., PF). DLG, dorsolateral geniculate nucleus; LD, laterodorsal nucleus; MD, mediodorsal nucleus; PF, parafscicular nucleus; Po, posterior thalamic nuclear group; STh, subthalamic nucleus; VM, ventromedial nucleus; VP, ventral posterior nucleus; ZIR, zona incerta rostral part.

***Calsyntenins*** encode transmembrane proteins with two extracellular cadherin domains [38]. They have recently been implicated in synaptogenesis through interactions with neurexins [39]. As with previous reports, we find Clstn1 to be broadly (but not ubiquitously) expressed across the thalamus [38,39], while Clstn2 is expressed more selectively (Figure 4; see also Figure 17, below), in contrast to report by Hintsch *et al.* of only background expression. Clstn3 mRNA was not detectable.

**Figure 4.**
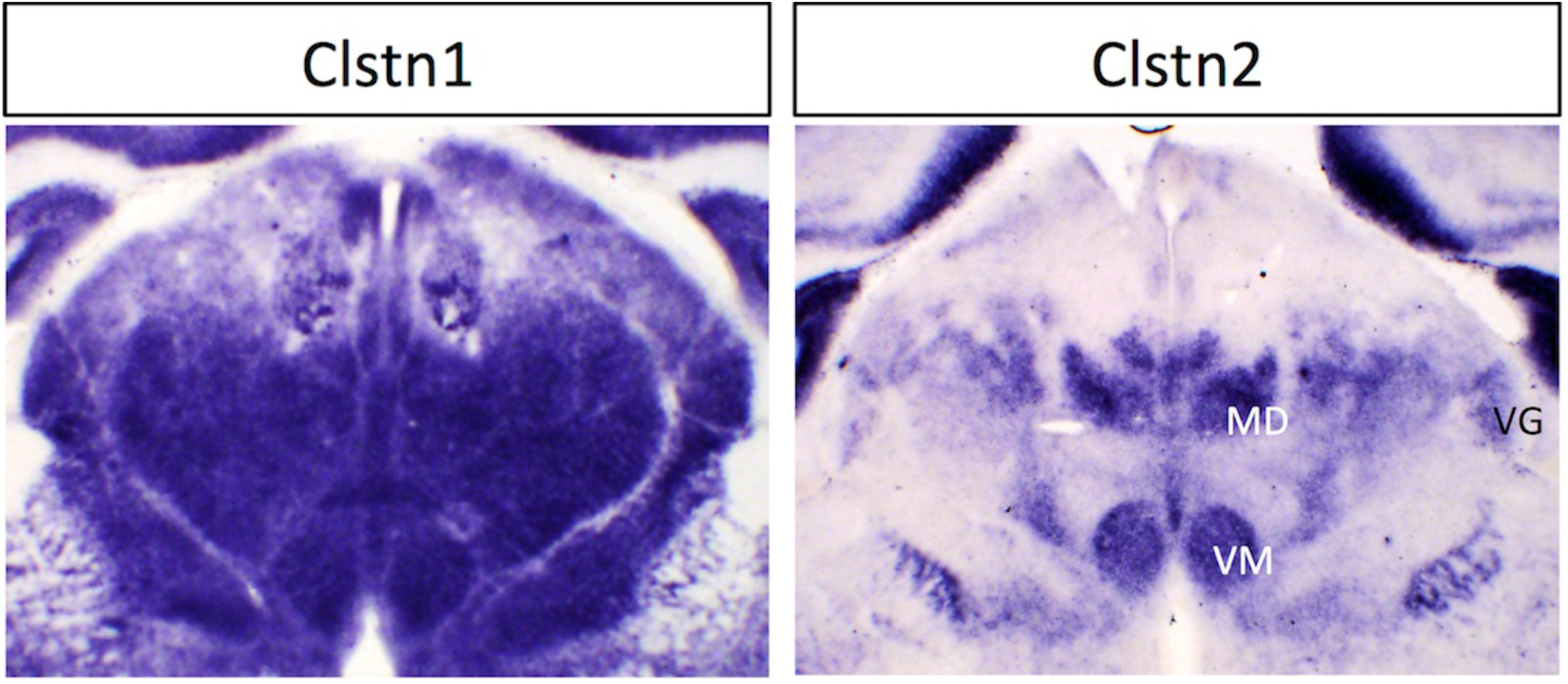
***In situ* hybridization patterns for Clstn family genes in dorsal thalamus at P0.** One coronal section for Clstn1 and Clstn2 is shown. Additional sections from the devABA are shown in Figure 17. MD, mediodorsal nucleus; VG, ventral geniculate nucleus; VM, ventromedial nucleus.

### Clustering analyses of developmental gene expression patterns in the thalamus

As a complementary strategy to identify candidate connectivity labels, we analysed data from the devABA, extending our search beyond genes conserved between vertebrates and invertebrates. The devABA provides qualitative and quantitative data on expression of 2002 genes based on mRNA *in situ* hybridization. These genes were chosen based on functional relevance to brain development or disorders of the brain. The expression density values for each gene in the atlas have been mapped to a three-dimensional digital template, with individual voxels attributed to different brain regions. We therefore extracted the expression density values for voxels attributed to the dorsal thalamus for 2002 genes at each of seven ages: embryonic day 11.5 (E11.5), E13.5, E15.5, E18.5, postnatal day 4 (P4), P14 and P28 (Table S2). These time-points are developmentally relevant to regional specification/patterning (E11.5), axon pathfinding (E13.5, E15.5 and E18.5), synaptogenesis (P4 and P14) and cortical plasticity (P28). Because we were more interested in comparing temporal profiles than relative levels across genes, we normalised these data for each gene by dividing by the average expression value for that gene across all ages (Table S3 Columns B-H).

In order to find genes with similar temporal expression profiles we used *k*-means clustering, which clusters elements of a matrix into a user-defined number of clusters (the *k* value). Since there is not necessarily a “correct” number of clusters, we performed *k*-means clustering with values of *k* from 6 to 18 and examined the flux in clustering outputs across these levels. As the input value of *k* increased, clusters tended to get subdivided as opposed to reshuffled, giving a roughly hierarchical organisation (Table S3 Columns I-U; Figure 5). We selected *k* = 10 for further data analysis, since the data segregated into clusters differentially peaking at all embryonic and postnatal time-points, while leaving large enough clusters for statistical analyses of enrichment (Figure 6). Lower *k* values gave poorer separation across the embryonic and early postnatal ages of highest interest to us, while higher values gave increasing refinement only at later postnatal stages (Figure S7).

**Figure 5.**
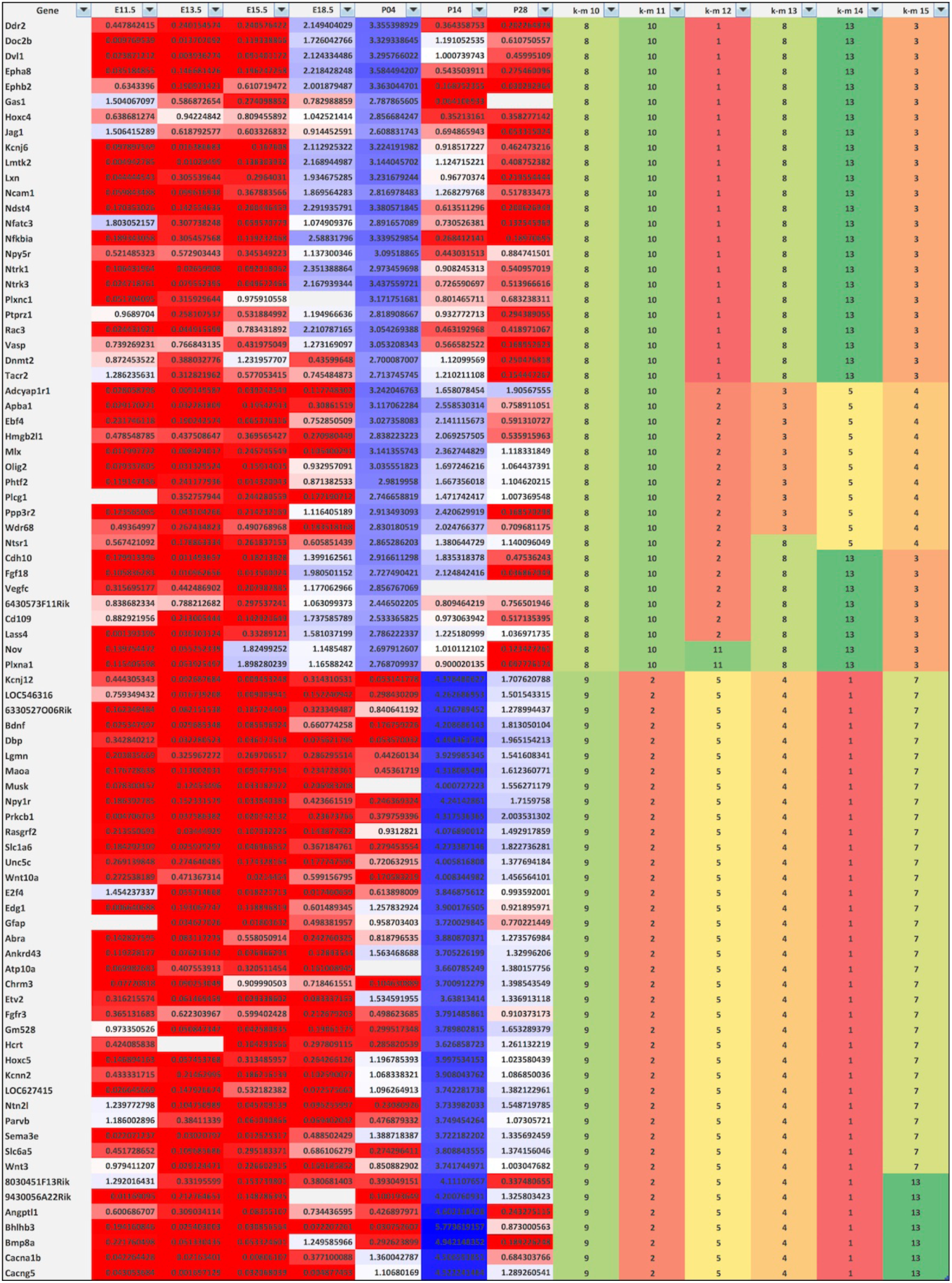
**Excerpt of *k*-means clustering analyses.** Clustering of a small subset of genes is shown to illustrate the method. Gene names are shown in column 1. Columns 2 to 8 show normalised expression densities per gene per time-point. Heatmap’s 3 colour scale of gene expression data: 0.2, red; 1, white; 5, blue. Columns 9 to 14 show cluster numbers assigned under *k* = 10 to *k* = 15 clustering, with clusters colour-coded to facilitate visualisation. Clusters are sorted at *k* = 10, followed by *k* = 11, followed by *k* = 12 and so on until *k* = 15. As the input value of *k* increases, clusters tend to get subdivided as opposed to reshuffled, giving a roughly hierarchical organisation. A clear difference in expression pattern between clusters 8 and 9 at *k* = 10 is evident, with cluster 8 showing peak expression at P4, and cluster 9 showing peak expression at P14. Within cluster 8, additional subclusters become apparent at *k* = 12 and above. The full clustering output for all 1996 genes is shown in Table S3.

**Figure 6.**
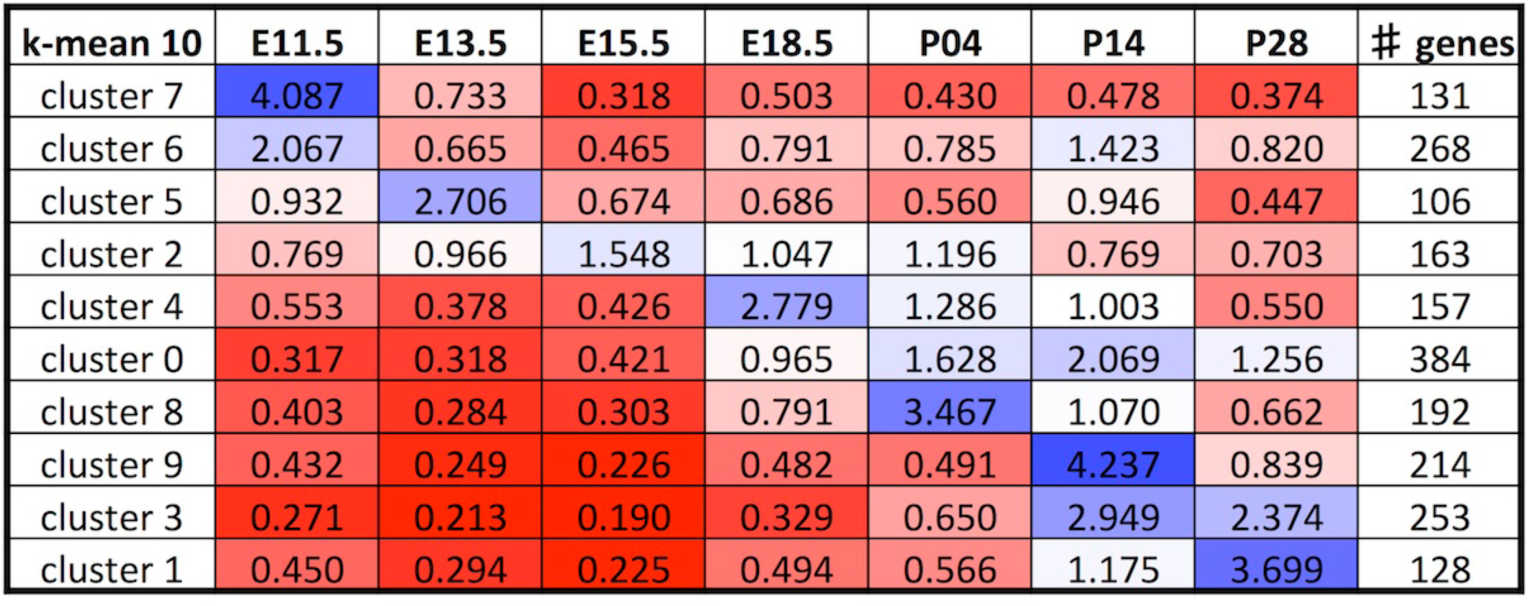
**Summary of expression profiles at *k* = 10.** Normalised expression densities were averaged per cluster to see the trends of expression at *k* = 10. Clusters were organised chronologically with those showing early peaks of expression at the top and later peaks of expression at the bottom. Heatmap’s 3 colour scale of gene expression data: 0.2, red; 1, white; 5, blue. Number of genes per cluster is shown in the rightmost column.

### Gene annotation and enrichment analyses

We used two different approaches to annotate genes, based on known or predicted localisation of the protein product or on known biological function. First, we screened protein products for cellular localisation with an emphasis on the extracellular environment by assessing the outputs of several software tools available online (PSORTII, SignalP, Big-PI Predictor and TMHMM; Table S4), along with manual curation based on literature sources. The devABA dataset predominantly comprises proteins found in the nucleus (907 proteins), followed by 293 secreted proteins, 286 GPI-anchored or single-pass transmembrane proteins, 267 cytoplasmic (or in organelles) proteins and 240 multi-pass transmembrane proteins (Table S5 Column K). Three genes could not be annotated: C030002O17Rik, LOC433436 and mCG146432.

Second, we grouped genes by ascribed functions, using GO term annotations and systematic manual curation (Table S5 Column M). The dataset was divided into 6 groups: Group 1: axon guidance pathway and cell adhesion; Group 2: synapse; Group 3: receptor tyrosine kinases and their ligands, and patterning; Group 4: neurotransmission pathway (G-protein-coupled receptors, ion channels, gap junctions); Group 5: chromatin and transcription factor activity; Group 6: other (cytoskeleton, extracellular matrix, myelin, metabolic enzymes and signal transduction) and unannotated.

When a gene fell into more than one group, an order of priority was used (the order in which the groups are presented); for example, a gene encoding a receptor tyrosine kinase that is involved in axon guidance would appear in Group 1 and not in Group 3. Our dataset predominantly included genes involved with chromatin and transcription factor activity (677 genes in Group 5); followed by 360 genes expressing receptor tyrosine kinases and their ligands, and genes involved in patterning (Group 3); 286 genes involved in the axon guidance pathway and cell adhesion (Group 1); 270 in the neurotransmission pathway (Group 4); and, 91 genes involved with synapses (Group 2; Table S5). Group 6 contained 107 genes involved with or part of the cytoskeleton, extracellular matrix, myelin, signal transduction pathways or expressed metabolic enzymes. This group also included 205 unannotated genes to bring the total to 312 genes.

We hypothesized that different types of proteins would be enriched in clusters with peak expression at time-points relevant to specific developmental functions. In order to test this, we compared the observed numbers for each category in each cluster to the expected value by normal distribution.

A variety of trends emerged from these data (Figures 7 and 8). Chi square statistics on the whole dataset showed deviation from expected by random distribution at p-value < 0.001 for either localisation or function (Table 2A and B). Further statistics were done on individual categories to establish which ones were enriched in particular clusters. The Bonferroni correction was taken into account due to multiple testing, lowering the threshold to p-value ≤ 0.01 and p-value ≤ 0.008 for localisation and function, respectively. For localisation, the trends observed with secreted and cytoplasmic proteins did not reach statistical significance (p-value > 0.1). However, the distribution of single-pass transmembrane and GPI-anchored proteins, multi-pass transmembrane proteins and nuclear proteins were significant at p-value < 0.0001, p-value < 0.001 and p-value < 0.001, respectively. For functional groups, all trends observed reached statistical significance, at p-value < 0.005 for Group 2, p-value <0.001 for Group 1, and p-value < 0.0001 for Groups 3 to 6.

**Table 2A.**
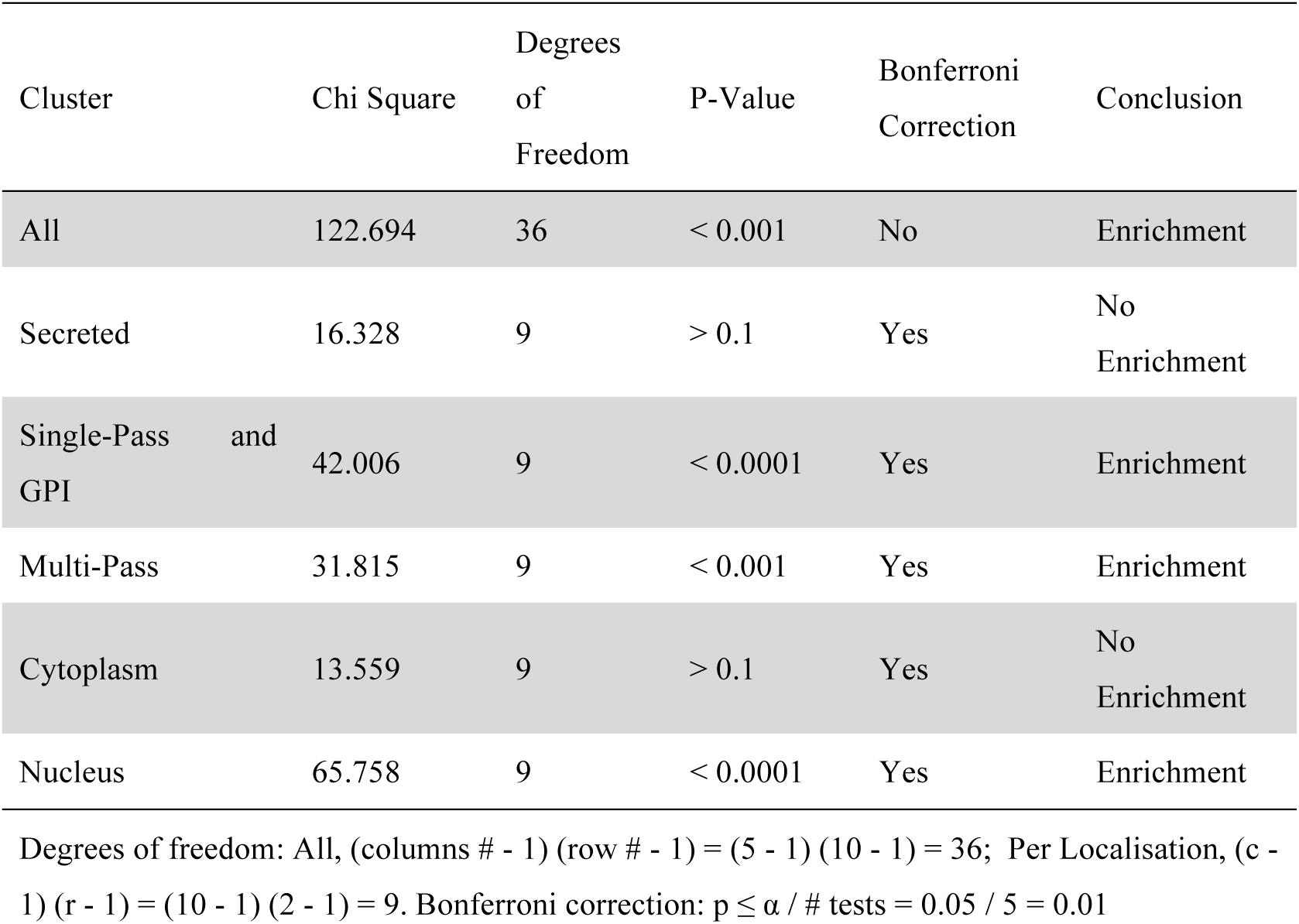
A Enrichment Analyses by Protein Localisation

**Table 2B.** Table 2B Enrichment Analyses by Functional Group

| Cluster | Chi Square | Degrees of Freedom | P-Value | Bonferroni Correction | Conclusion |
| --- | --- | --- | --- | --- | --- |
| All | 206.160 | 45 | $< 0.001$ | No | Enrichment |
| Group 1 | 30.770 | 9 | $< 0.001$ | Yes | Enrichment |
| Group 2 | 26.554 | 9 | $< 0.005$ | Yes | Enrichment |
| Group 3 | 46.368 | 9 | $< 0.0001$ | Yes | Enrichment |
| Group 4 | 41.989 | 9 | $< 0.0001$ | Yes | Enrichment |
| Group 5 | 73.622 | 9 | $< 0.0001$ | Yes | Enrichment |
| Group 6 | 37.325 | 9 | $< 0.0001$ | Yes | Enrichment |

Degrees of freedom: All, (columns # - 1) (row # - 1) = (6 - 1) (10 - 1) = 45; Per Group, (c - 1) (r -

By plotting the relative enrichment for each category across clusters, ordered by time-point of highest expression, it was possible to discern what relationships were driving these deviations from a random distribution (Figures 7 and 8), and to assess their biological plausibility. For the localisation data, nuclear proteins were enriched in clusters peaking at early time-points, whereas single-pass transmembrane and GPI-anchored proteins were enriched in clusters peaking from E15.5 to P4, and multi-pass transmembrane proteins in clusters peaking from P4 to P28 (Figure 7). This nicely correlates with the extensive requirement of transcription factors during patterning and differentiation early on (localised in the nucleus), followed by axon guidance molecules expressed extracellularly during axon pathfinding, and finally synaptogenesis and neurotransmission postnatally (including many multi-pass neurotransmitter receptors and ion channels). These inferences were supported by enrichment of functional groups: chromatin and transcription factors early on, axon guidance and synapse molecules in later embryogenesis, and neurotransmission pathway genes at later postnatal stages (Figure 8).

**Figure 7.**
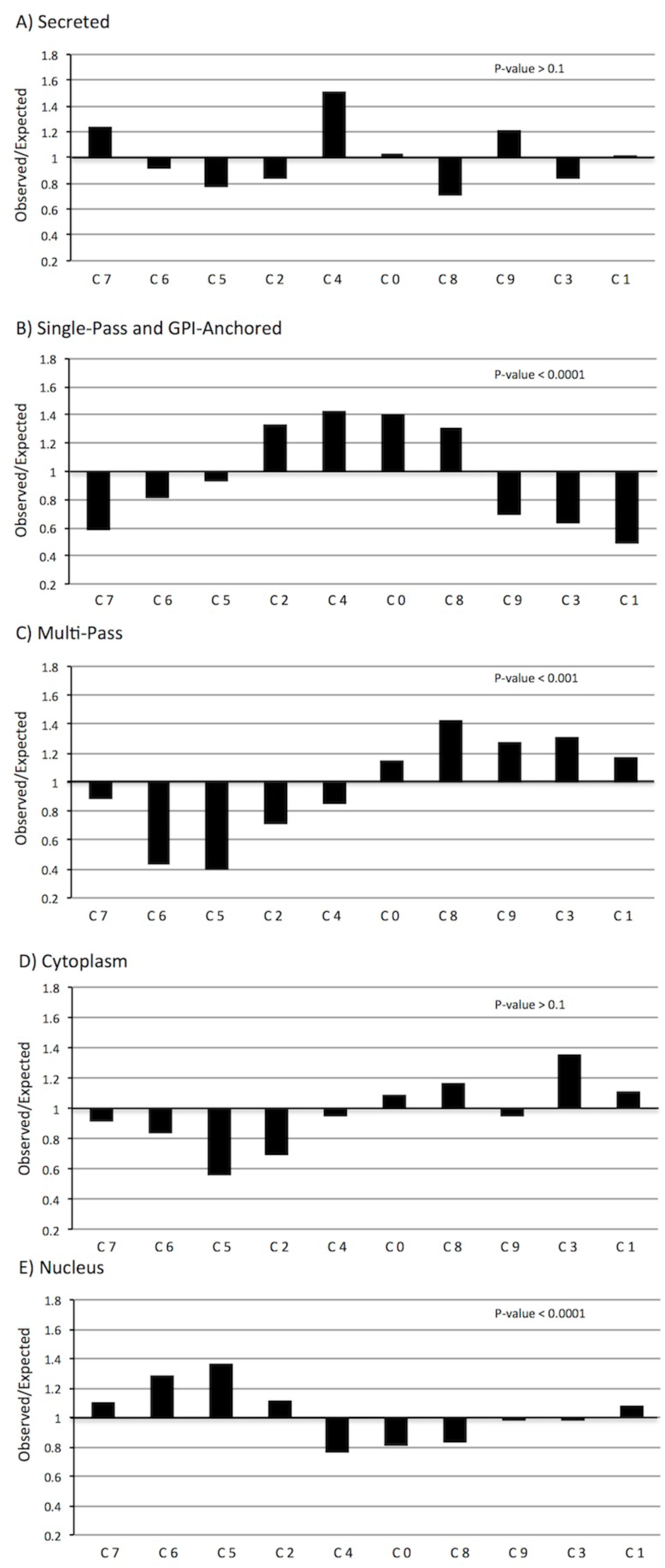
**Enrichment of protein localisation categories across clusters.** Relative enrichment across clusters is shown by plotting observed/expected numbers of proteins per cluster for each of five groups: secreted proteins (A), single-pass transmembrane or GPI-anchored proteins (B), multi-pass transmembrane proteins (C), cytoplasmic proteins (D) and nuclear proteins (E). A value of 1 indicates observed data matches expected values, whereas a value below or above 1 indicates decreased or increased counts compared to expected values, respectively. The ten clusters (C0-C9) are organized according to age of peak expression, as in Figure 6. P-values from chi square analyses are shown in upper right corner of each graph. Genes encoding nuclear proteins are enriched in clusters defined by strong early embryonic expression, single-pass transmembrane and GPI-linked protein-encoding genes are enriched in clusters expressed at mid-embryonic stages and multi-pass transmembrane proteins are enriched in clusters expressed at postnatal stages.

**Figure 8.**
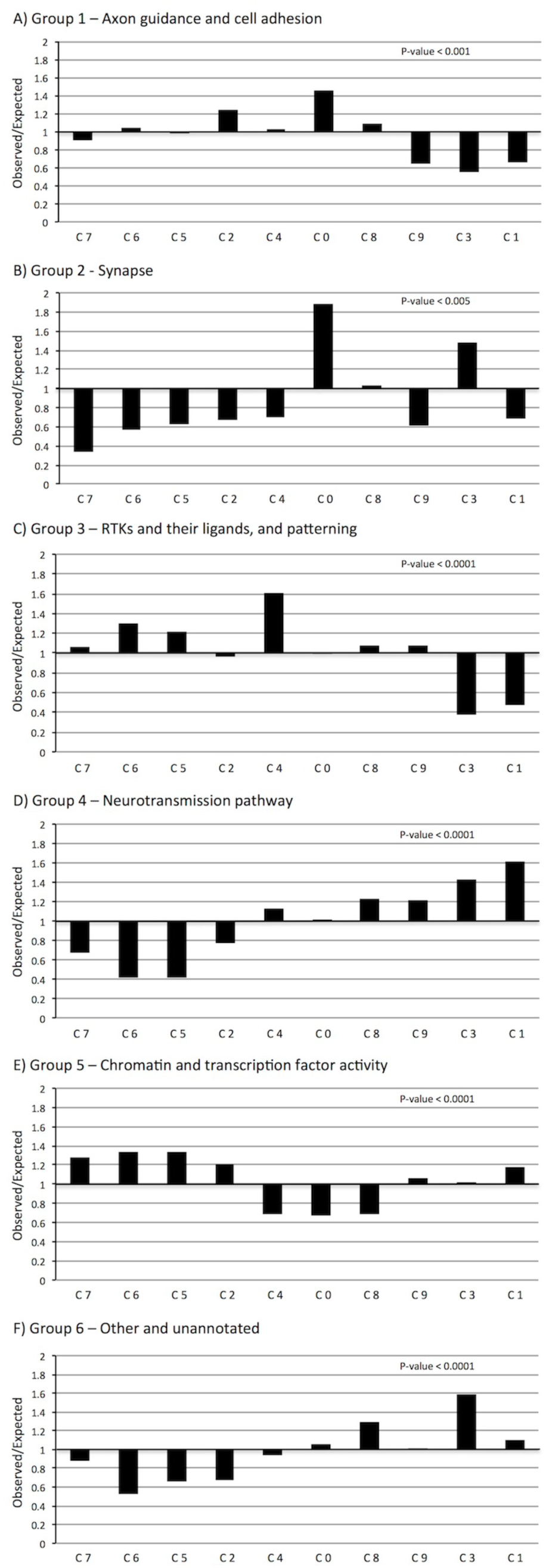
**Enrichment of functional groups across clusters.** Relative enrichment across clusters is shown by plotting observed/expected numbers of proteins per cluster for each of six groups: Group 1 - axon guidance pathway and cell adhesion (A); Group 2 - synapse (B); Group 3 - receptor tyrosine kinases and their ligands, and patterning (C); Group 4 - neurotransmission pathway (GPCRs, ion channels, gap junctions; D); Group 5 - chromatin and transcription factor activity (E); and, Group 6 - other (cytoskeleton, extracellular matrix, myelin, metabolic enzymes and signal transduction) and unannotated (F). A value of 1 indicates observed data matches expected values, whereas a value below or above 1 indicates decreased or increased counts compared to expected values, respectively. Clusters 0-9 are organized chronologically. P-values from chi square analyses are shown in upper right corner of each graph. All groups showed statistically significant deviation from expected distributions.

Taken together, these protein localisation and functional enrichment analyses provide a powerful and independent validation of our clustering analyses, indicating that the clusters we have identified with *k* = 10 track valid biological trends. They further suggest that clusters enriched for known molecules involved in axon guidance or synaptic connectivity may also contain novel molecules with these functions. In order to specifically search for such genes, we focused on clusters 2, 4, 0 and 8, which show enrichment for single-pass and GPI-linked proteins and for proteins involved in axon guidance or synaptogenesis. These have peak expression at E15.5 (cluster 2), E18.5 (cluster 4) and P4 (clusters 0 and 8).

### Identifying candidate connectivity labels

Clusters 2, 4, 0 and 8 had a total count of 896 genes of which 426 encode extracellular proteins. We screened through them by visual inspection of the devABA expression data at their peak expression to identify those with selective or differential expression across the thalamus. Genes showing uniform expression were not considered further. We generated a list of 215 genes that potentially encode specific connectivity information (Table S6). This dataset includes numerous genes already implicated in thalamocortical connectivity (such as Chl1, DCC, L1CAM, NCAM1, Ntn1, Robo1 and Robo2), which we did not characterize further. However, where previously implicated genes were part of larger families (e.g., cadherins, Ephrins and Eph-receptors, semaphorins), they were included for comparison with other members. Some genes had previously demonstrated roles in axon guidance or synapse formation in other contexts and were of high interest to us, as were those encoding surface proteins not previously implicated in these processes. However, we did not pursue any of the large number of neurotransmitter receptor and ion channel genes, given a lower prior probability of involvement as direct axon guidance or synaptic connectivity labels.

In order to present a summary of the expression patterns of these genes and to enable comparison across genes, we extracted images for each gene from the devABA database of a consistent lateral and/or a more medial or parasagittal section at E18.5 (Figure S8 and Figures 9–20). As the borders between nuclei are still developing at E18.5, it is not possible to definitively match expression patterns to specific nuclei at this stage. These figures are not intended to be comprehensive, as the full dataset can be viewed on the devABA website. They do, however, allow a survey of the general trends of expression patterns across the thalamus and a comparison of multiple members of specific gene families, while also highlighting numerous individual genes with strikingly selective patterns.

In general, the expression patterns were extremely diverse – in fact, we did not detect any two patterns that were identical across these genes. Some genes showed highly selective expression patterns, on in some developing nuclei and off in others. Others showed differential expression, higher in some nuclei than others, with substantial variation in extent of expression across the thalamus. Within developing nuclei, some genes appeared to be expressed at uniform levels, while others showed an uneven or graded distribution. Below, we consider the expression patterns of 73 genes organised by protein family or general function (Table 1 Columns B-C).

***Ephrins and Eph-receptors:*** Twelve members of the Eph and Ephrin gene families show differential or selective expression across the developing thalamus (Figure 9). Multiple members of the EphA and -B receptor and Ephrin-A and -B families have been implicated in diverse aspects of thalamic connectivity. The most extensively investigated have been EphA4, EphA7, EphrinA2 (Efna2) and EphrinA5 (Efna5). Roles for these genes have been demonstrated in parcellation of thalamic nuclei [19], target selection and topography of retinal axons into the thalamus [40], topographic guidance of thalamic axons through the ventral telencephalon [41], areal and layer-specific targeting and topography [reviewed in 42], reciprocal corticothalamic axon guidance and target selection [43,44] and even influences on cortical progenitor proliferation and differentiation dynamics [45]. EphB1 and -B2 have also recently been shown to mediate thalamic axon guidance through the ventral telencephalon [46]. The selective expression of several other members in the thalamus, including EphA3, -A5, -A6 and -A8 has also previously been noted [20,47]. The ABA data confirm these findings and allow a direct comparison of expression patterns across members of these families. In addition, we find that EphA1, EphA10, EphB6 and Efnb3 are also differentially expressed across the developing thalamus.

**Figure 9.**
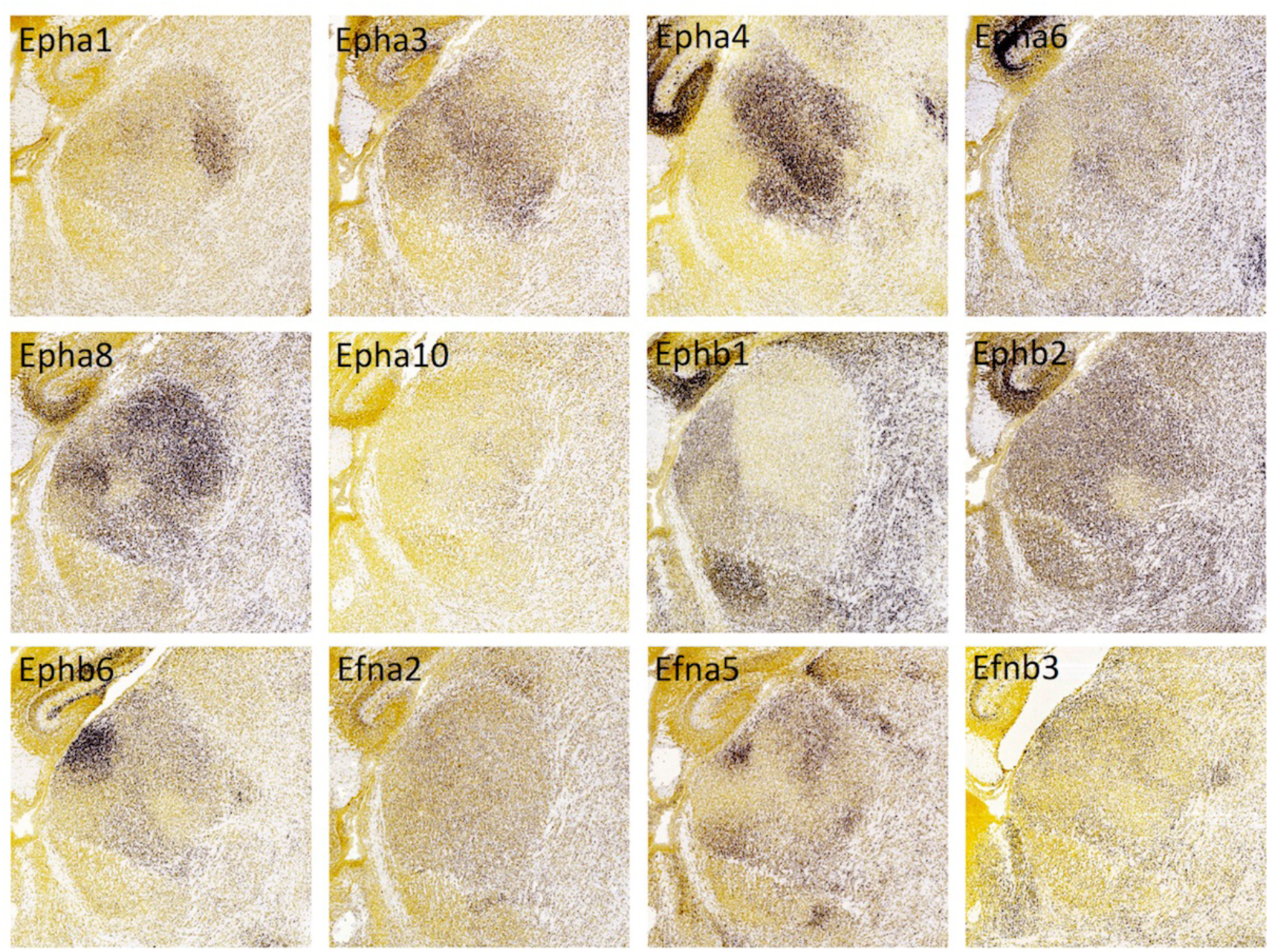
**Ephrin and Eph-receptor expression in the thalamus at E18.5.** *In situ* hybridization data from sagittal sections of the thalamus at E18.5 obtained from the devABA revealed differential patterns of expression of Eph-receptor (Eph) A1, A3, A4, A6, A8, A10, B1, B2, B6, and Ephrin (Efn) A2, A5 and B3. See text for details. A representative and equivalent section of medial thalamus is shown for each gene (see Figure S7 for details on how the sections were selected).

***Cadherins:*** We report differential or selective expression of twelve members of the cadherin family across the developing thalamus (Figure 10). A number of cadherins have been implicated previously in thalamic connectivity. Takeichi and colleagues showed differential expression of Cdh6, Cdh8 and Cdh11 in both cortex and thalamus and used dye-tracing to demonstrate that Cdh6- or Cdh8-positive cortical areas are specifically connected with Cdh6- or Cdh8-positive thalamic nuclei, respectively [48]. Similarly, but for afferent connections to the thalamus, Cdh6 has been shown to act as a homophilic targeting molecule for a subset of retinal ganglion cells which project to nuclei mediating non-image-forming visual functions [10]. N-Cadherin (Cdh2) is required for proper termination of thalamic axons in layer 4 of the cortex [49], and Cdh2 and Cdh8 differentially label thalamic terminations in somatosensory cortex barrel center and septal compartments, respectively [50]. Differential expression across thalamic nuclei of the remaining cadherins in this group, Cdh4, 7, 9, 10, 12, 13, 24 and the atypical cadherin Fat3, has not, to our knowledge, been previously reported and greatly expands the possible involvement of this gene family in specifying thalamic connectivity. The diversity of expression patterns is particularly noteworthy; even in this single parasagittal section, no two members of this family show an identical pattern. Given the known potential for heteromeric complex formation between cadherins, these overlapping patterns generate a very large potential combinatorial code.

**Figure 10.**
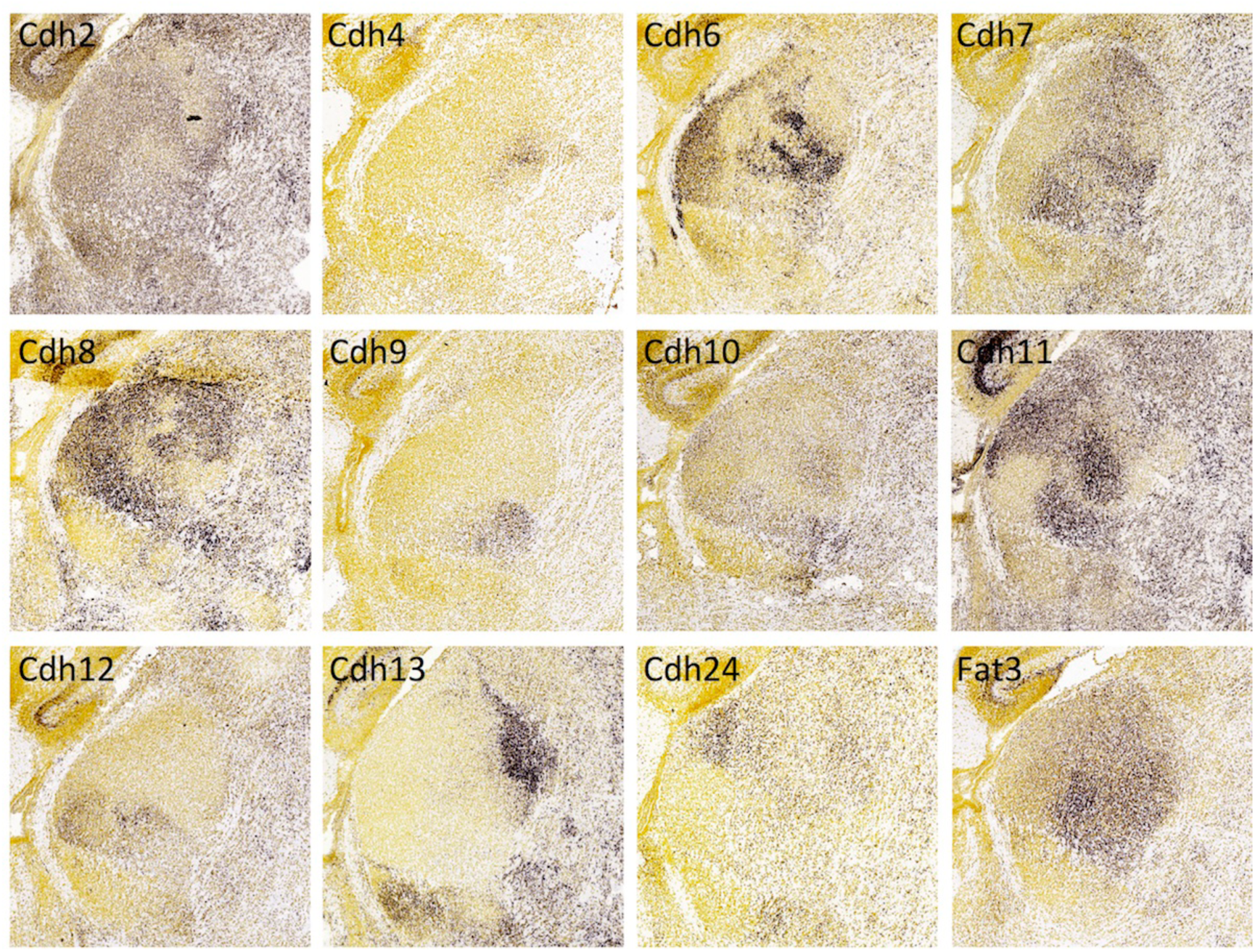
**Cadherin expression in the thalamus at E18.5**. *In situ* hybridization data from sagittal sections of the thalamus at E18.5 obtained from the devABA revealed differential patterns of expression of Cadherin (Cdh) 2, 4, 6, 7, 8, 9, 10, 11, 12, 13, 24 and Fat3. See text for details. A representative and equivalent section of medial thalamus is shown for each cadherin (see Figure S7 for details on how the sections were selected).

***Protocadherins***: A number of members of the unclustered protocadherin family show differential expression across the thalamus at E18.5, including Pcdh1, 10, 11X, 19 and 21 (Figure 11). We note that Pcdh18 also shows differential thalamic expression at this stage (not shown) but is not included in our clusters of interest due to an earlier time-point of peak expression. These results replicate previous findings reported by Kim et al (2007), who also noted differential expression across thalamus of additional members of this family that are not in the devABA dataset (Pcdh7, 8, 9, 15 and 17) [51]. The functional importance of these genes in thalamic connectivity has not been directly tested. Though defects in thalamocortical axon projections occur in Pcdh10^-/-^ mice, they have been attributed to functions in patterning of the ventral telencephalon and not to activity in thalamic axons themselves [52]. Many of the protocadherins are also expressed in selective patterns across cortical areas and layers [51], suggesting a possible interaction code between interconnected regions.

**Figure 11.**
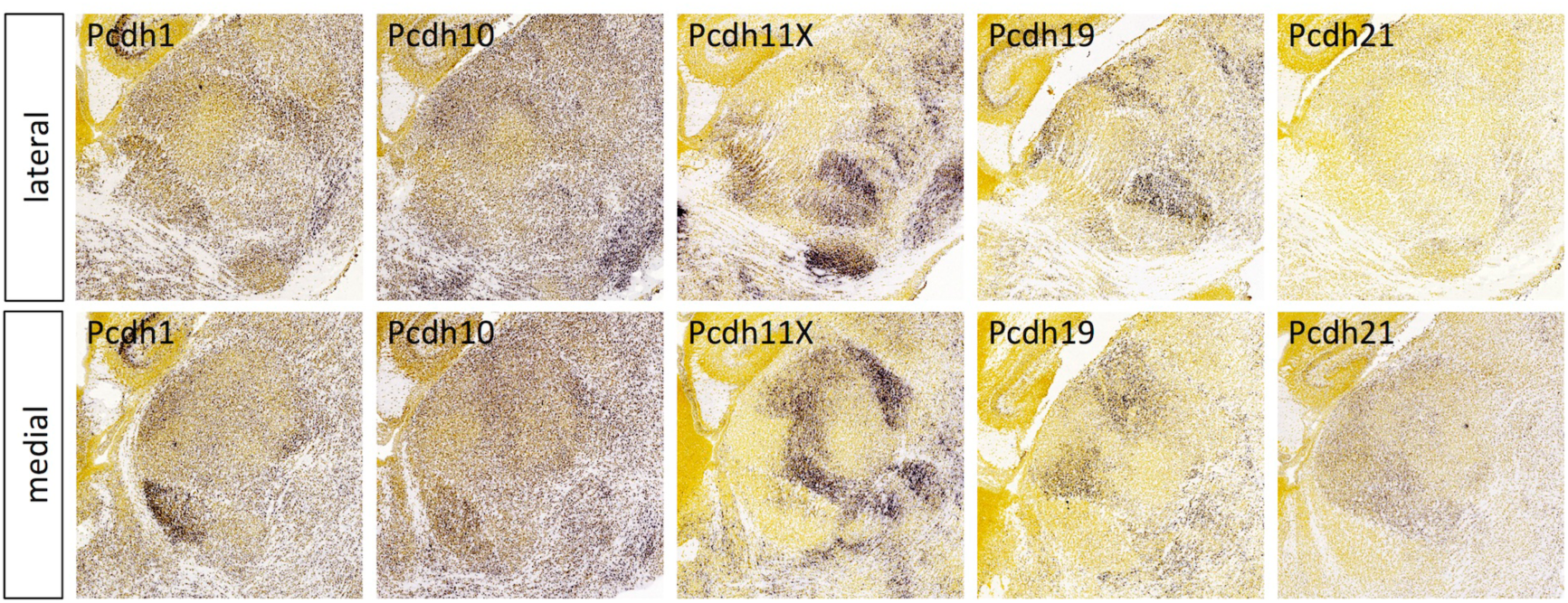
**Protocadherin expression in the thalamus at E18.5**. *In situ* hybridization data from sagittal sections of the thalamus at E18.5 obtained from the devABA revealed differential patterns of expression of Protocadherin (Pcdh) 1, 10, 11X, 19 and 21. See text for details. Two representative and equivalent sections are shown for each protocadherin, one lateral and the other medial (see Figure S7 for details on how the sections were selected).

***Semaphorins and Plexins:*** Among the members of the semaphorin and plexin gene families in the devABA dataset, the ones showing the most selective or differential expression across the thalamus are: Sema3F, Sema6A, Sema7A, PlxnA1, PlxnA2 and PlxnC1 (Figures 12, 13). PlxnB1 and Sema6D are also expressed in developing thalamus, but weakly and more uniformly (not shown). Sema3F has been previously implicated in thalamic axon guidance through the ventral telencephalon, and Sema7A in regulating thalamocortical axon branching in layer 4 of cortex [53,54]. However, these functions depend on expression of the semaphorins in these other regions, not in the thalamus itself, where differential expression has not been previously reported. Null mutants of *Sema6A* show misrouted thalamocortical axons [55], specifically from the dLGN [56]; this function may depend on expression of Sema6A in the thalamus itself or in the ventral telencephalon. No roles in thalamic axon guidance or connectivity have yet been described for PlxnA1, PlxnA2 or PlxnC1. The expression of Sema7A and its receptor PlxnC1 are notably complementary, while Sema6A and its binding partner PlxnA2 are expressed in an overlapping fashion, with Sema6A more widespread and PlxnA2 restricted to medial regions. Neuropilins also act as co-receptors for secreted semaphorins and have been implicated in thalamic axon guidance in response to Sema3F, signaling through Neuropilin-2-NrCAM co-receptor complexes [57] or Sema3A, acting through Neuropilin-1-CHL1 co-receptors [58]. Npn2 is not in the ABA developmental dataset but Npn1 is found in cluster 4 and is differentially expressed across thalamic nuclei, as previously described (not shown).

**Figure 12.**
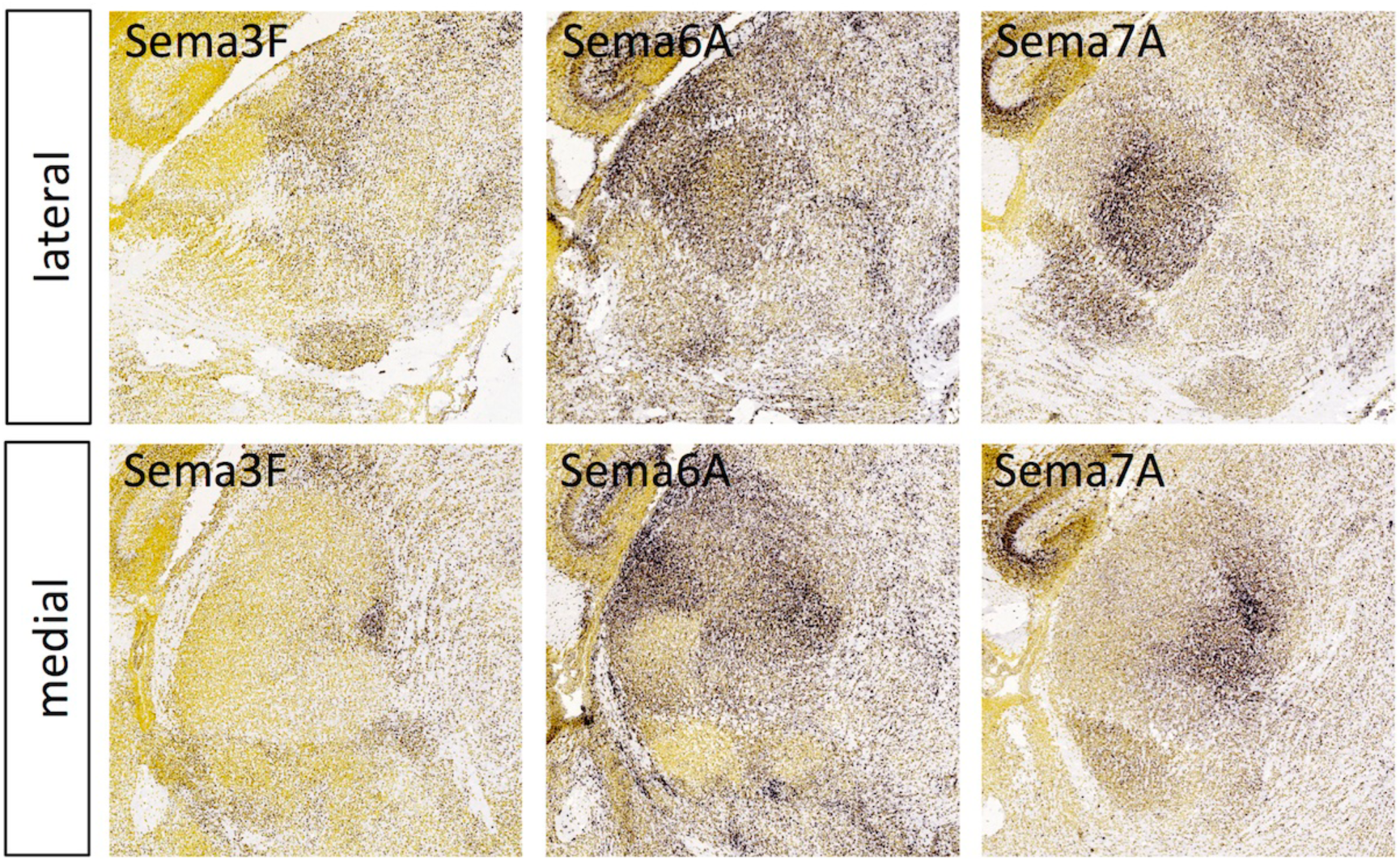
**Semaphorin expression in the thalamus at E18.5**. *In situ* hybridization data from sagittal sections of the thalamus at E18.5 obtained from the devABA revealed differential patterns of expression of semaphorins Sema3F, Sema6A and Sema7A. See text for details. Two representative and equivalent sections are shown for each semaphorin, one lateral and the other medial (see Figure S7 for details on how the sections were selected).

**Figure 13.**
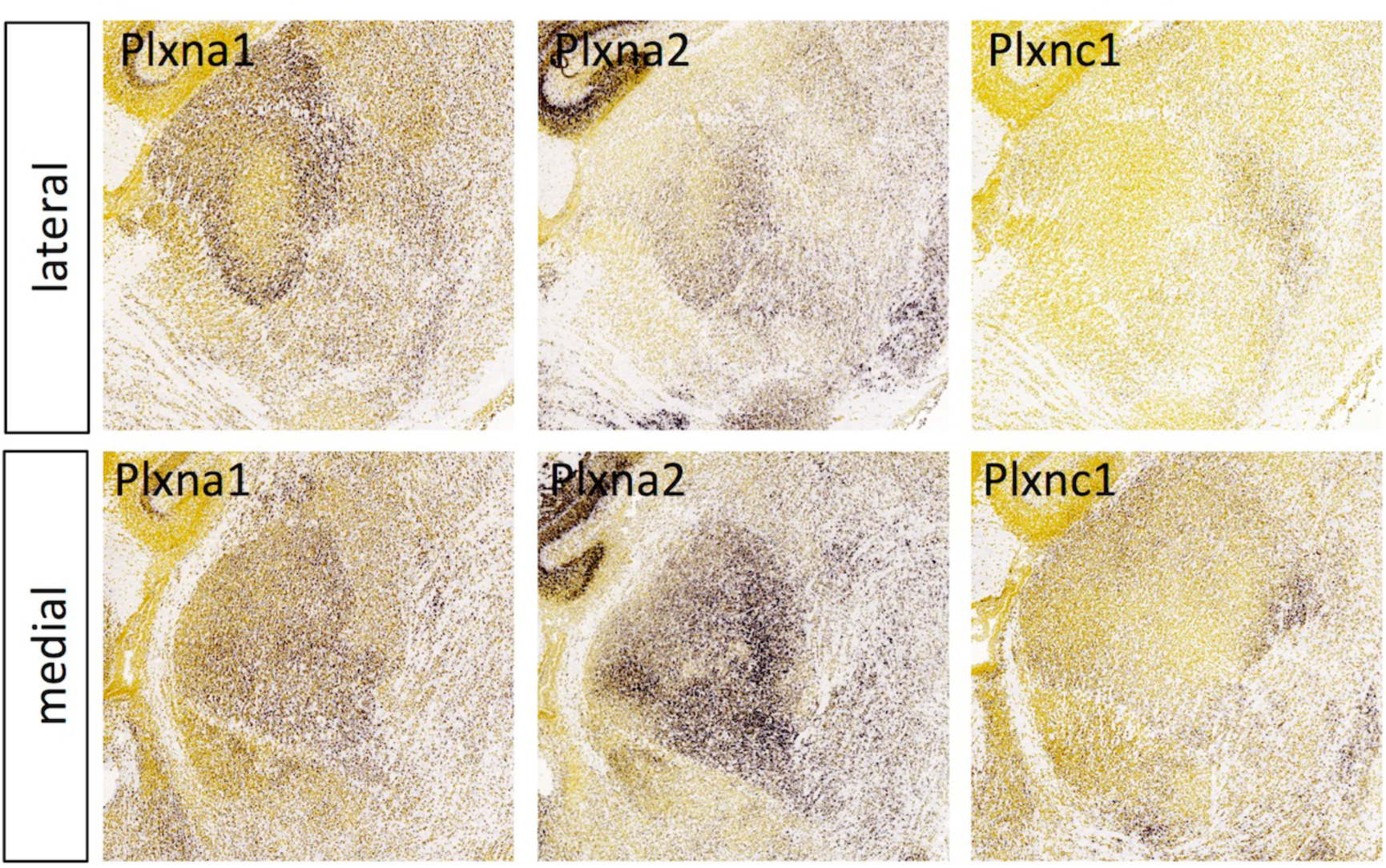
**Plexin expression in the thalamus at E18.5**. *In situ* hybridization data from sagittal sections of the thalamus at E18.5 obtained from the devABA revealed differential patterns of expression of plexins PlxnA1, PlxnA2 and PlxnC1. See text for details. Two representative and equivalent sections are shown for each plexin, one lateral and the other medial (see Figure S7 for details on how the sections were selected).

***Receptor tyrosine kinases and phosphatases:*** In addition to the Eph family of receptor tyrosine kinases, we identified a number of other members of this superfamily and one receptor tyrosine phosphatase showing quite selective expression in the thalamus (Figure 14). Kinases Flt3, Kit (cKit) and Ret have well-documented functions in the immune system and in various cancers, while their functions in the nervous system have been explored to varying extents. No functions for Flt3 in the nervous system have been reported. Kit has been shown to have effects on cortical neuronal migration and axonal extension [59], but has not been functionally implicated in thalamic development, though Kit-ligand (Steel) is also highly expressed in developing thalamus [20]. Ret has multiple, well-described roles in development of the enteric nervous system, as a receptor for GDNF, and mutations in this gene are an important cause of Hirschsprung disease. While expression in thalamus has been noted before [60], this has not been described in detail nor has a functional role in thalamic development been established. The neurotrophin receptors Ntrk2 (TrkB) and Ntrk3 (TrkC) have been studied in this context, with a demonstration that TrkB in particular is required for normal segregation of thalamic afferents in barrel cortex [61]. Ptpru (also called RPTP-lambda or PTP-RO) is closely related to RPTP-kappa and RPTP-mu and similarly mediates homotypic adhesion [62]. Ptpru has also been shown to associate with c-Kit and to negatively regulate its signaling [63]; however, the expression of these genes in thalamus appears largely non-overlapping (Figure 14). There are no published reports of Ptpru function in the nervous system, though many other members of the RPTP family play important roles in axon guidance and synaptic connectivity [64].

**Figure 14.**
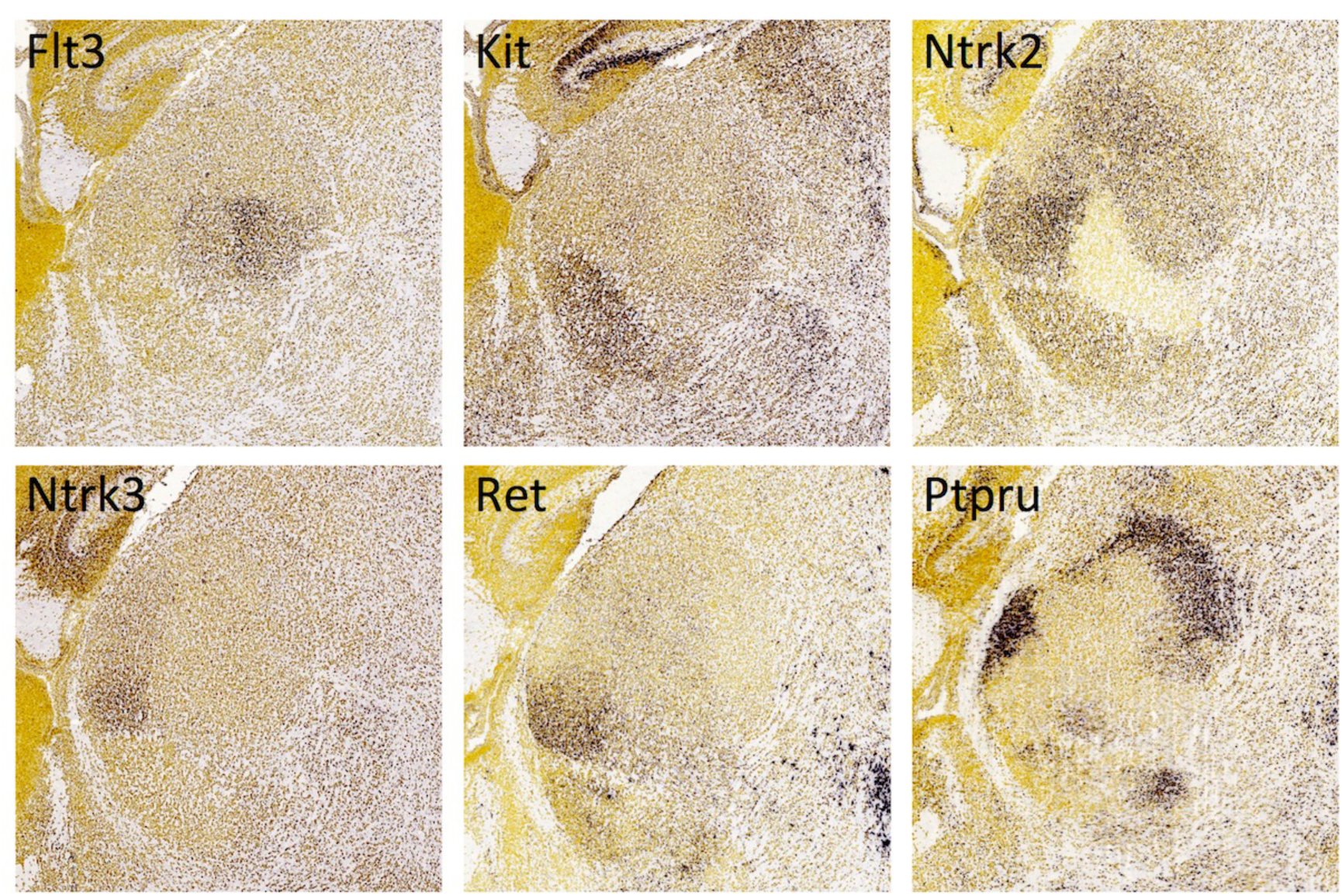
**Expression of receptor tyrosine kinases and phosphatases in the thalamus at E18.5**. *In situ* hybridization data from sagittal sections of the thalamus at E18.5 obtained from the devABA revealed differential patterns of expression of Flt3, Kit, Ntrk2, Ntrk3, Ret and Ptpru. See text for details. A representative and equivalent section of medial thalamus is shown for each gene (see Figure S7 for details on how the sections were selected).

***Immunoglobulin superfamily.*** Multiple Ig superfamily molecules have been implicated in axon guidance or neuronal connectivity in the developing thalamus, including Robo proteins, DCC, various cell adhesion molecules (NCAM, L1, CHL1…) and others [reviewed in 9]. Here, we highlight the selective expression of five Ig superfamily members not previously implicated in thalamocortical development (Figure 15). Alcam (also known as CD166, Neurolin, or DM-GRASP/SC1/BEN) plays a role in guidance and fasciculation of motor and retinal axons [65], and in topographic mapping of retinal axons across the superior colliculus, possibly through interaction with adhesion molecule L1 on retinal ganglion cells [66]. Cadm1 (better known as SynCAM) is involved in synapse organisation [67] and axon guidance [68]. Cd47 is a multi-pass transmembrane molecule with a single extracellular Ig domain. It acts as a receptor for presynaptic organising molecule SIRPalpha [69] and has been implicated in various processes in cerebellar development [70]. Cntn6 (NB3) is a member of the contactin family of adhesion molecules, which have widespread roles in neural development [71]. *Cntn6* null mutants have defects in synapse formation in cerebellum [72] and hippocampus [73]. Our results replicate previous reports of expression of Cntn6 in anterior thalamic nuclei [74,75]. Mdga1 is expressed in a layer- and area-specific manner in the cortex [76] and mutation of the gene leads to a delay in cortical neuronal migration [77]. More recently, Mdga1 has also been found to interact with neuroligin-2 to negatively regulate inhibitory synapse formation [78,79].

**Figure 15.**
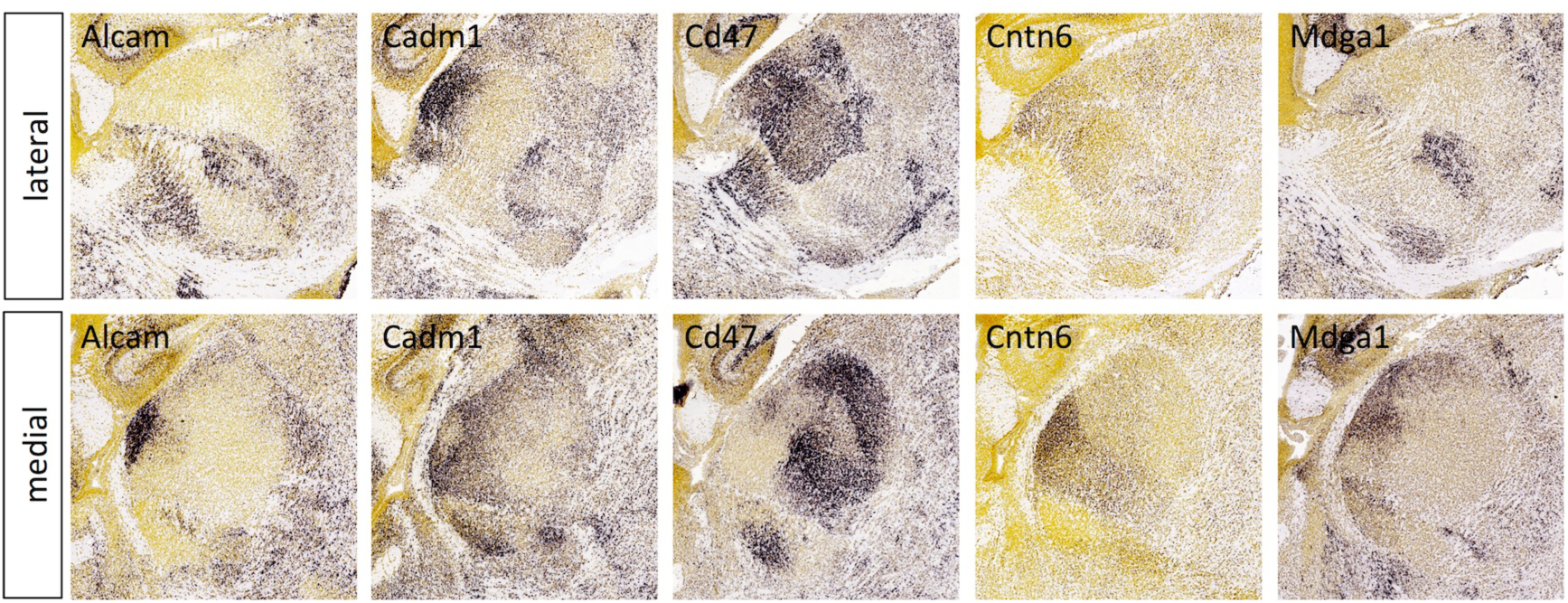
**Expression of immunoglobulin superfamily molecules in the thalamus at E18.5**. *In situ* hybridization data from sagittal sections of the thalamus at E18.5 obtained from the devABA revealed differential patterns of expression of Alcam, Cadm1, Cd47, Cntn6 and Mdga1. See text for details. Two representative and equivalent sections are shown for each gene, one lateral and the other medial (see Figure S7 for details on how the sections were selected).

***Leucine-rich repeat superfamily:*** Many members of the extracellular leucine-rich repeat (LRR) superfamily have been implicated in axon guidance, synaptic target selection and other aspects of neural development [23,80]. LRR superfamily genes are highly under-represented in the devABA dataset, with no members of the Lrfn, Slitrk, NGL, Elfn or LINGO subfamilies and only one representative of each of the Lrrtm, Lrrc, Lrrn and Amigo subfamilies. Amongst the LRR genes that are represented, we identified three with selective expression in developing thalamus: Lgi2, Lrrn3 and Rtn4rl1 (Figure 16). Lgi2 encodes a secreted LRR protein of the Lgi subfamily that have been implicated in synaptic development and epilepsy [81]. Lgi1 regulates synapse formation and function through interactions with postsynaptic Adam22 and presynaptic Adam23 proteins [82,83]. Lgi2 likely has similar functions as it also interacts with these Adam proteins and mutations in the gene cause epilepsy in dogs [84]. Lrrn3 (or Nlrr3) is a member of the LRR-Ig-FN3 group [23]. Roles for Lrrn1 and Lrrn2 have been described in hindbrain development in chick [85,86]; otherwise this family remains poorly characterised. Selective expression in dorsal thalamus has been commented on previously, in particular its complementarity to the expression pattern of Slitrk6 [87]. Rtn4rl1 is also known as Nogo-receptor-3, NgR3. NgR’s were first isolated as receptors for the inhibitory myelin protein Nogo-66, but also function as receptors for chondroitin proteoglycans [88] and have been implicated in synaptic development in hippocampus [89].

**Figure 16.**
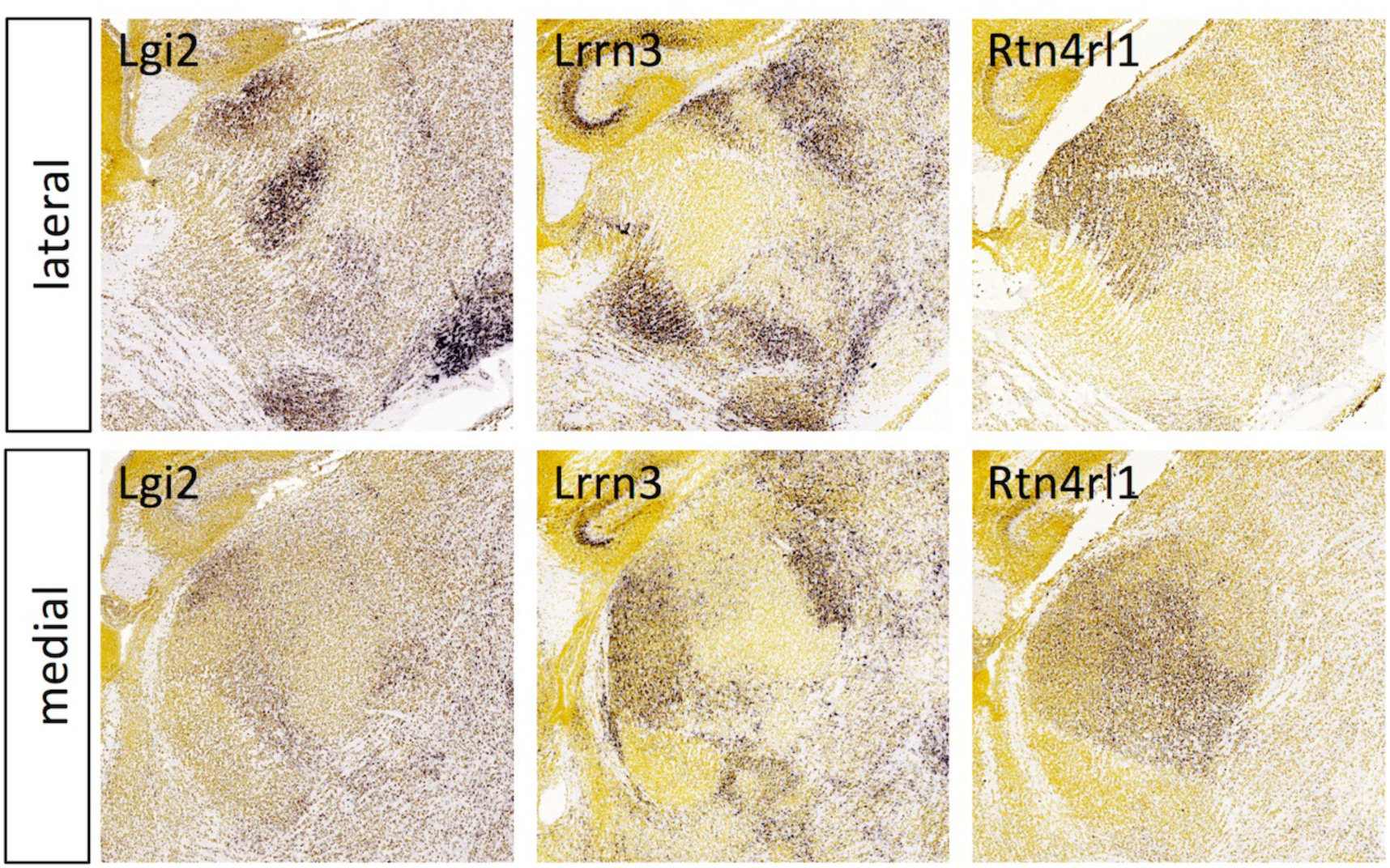
**Expression of leucine-rich repeat superfamily members in the thalamus at E18.5**. *In situ* hybridization data from sagittal sections of the thalamus at E18.5 obtained from the devABA revealed differential patterns of expression of Lgi2, Lrrn3 and Rtn4rl1. See text for details. Two representative and equivalent sections are shown for each gene, one lateral and the other medial (see Figure S7 for details on how the sections were selected).

***Small gene families***: A number of small gene families were identified with compelling expression patterns in developing thalamus (Figures 17, 18).

***Calsyntenins***: The devABA data replicate our own *in situ* hybridization results (Figure 17), with widespread, but not ubiquitous expression of Clstn1 and much more selective expression of Clstn2.

***Cerebellins*** encode secreted molecules with synaptogenic activity. They interact with multiple partners including the glutamate receptor subunits GluRδ2 and neurexins [90] as well as the netrin receptor DCC [91]. We find strong and selective expression of Cbln2 and Cbln4 across thalamic nuclei (Figure 17). Their differential expression across the brain has been documented previously, including in the developing thalamus [92]. By contrast, Cbln3 is not expressed outside the cerebellum. Cbln1 is not in the devABA dataset but is also expressed in subsets of thalamic nuclei [92].

**Figure 17.**
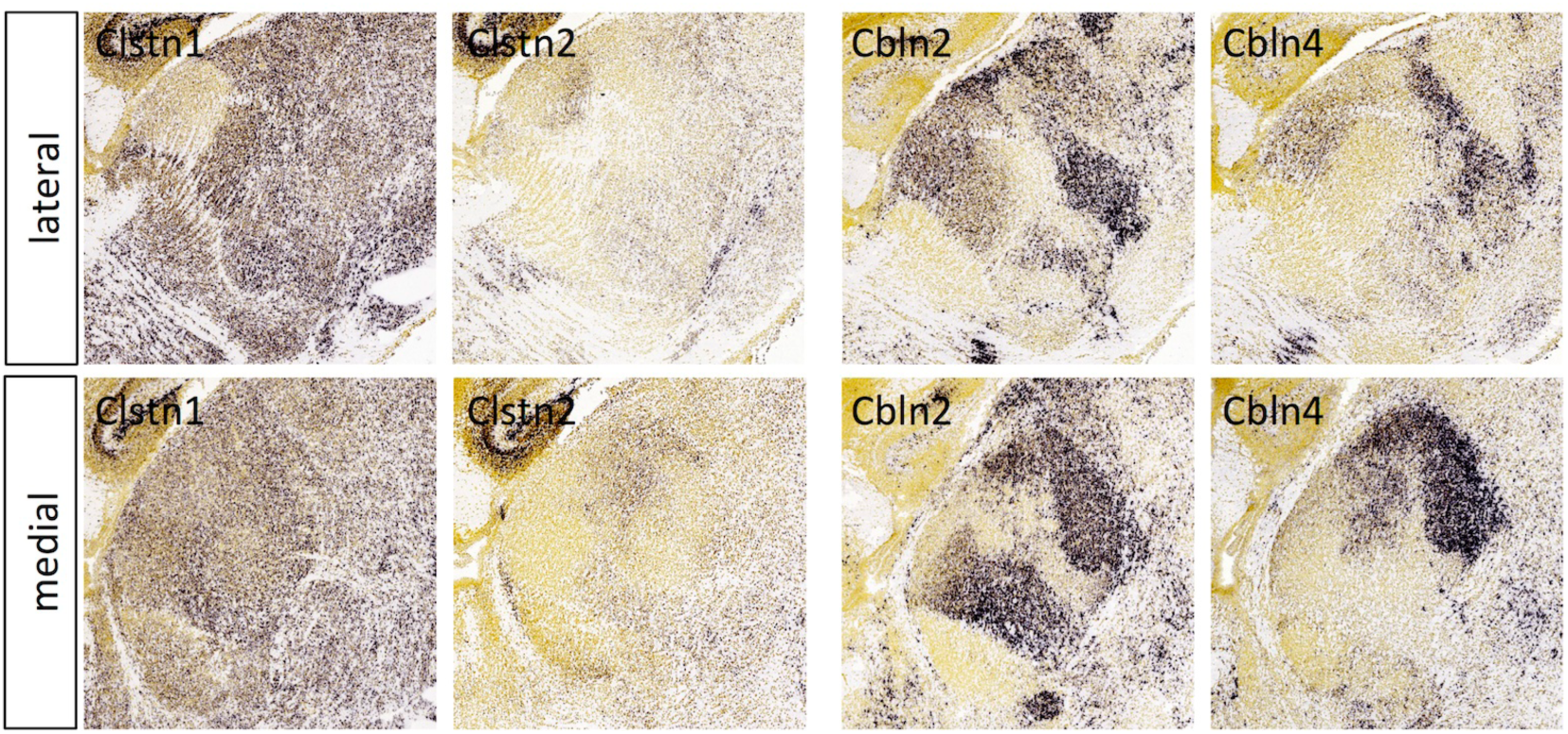
**Calsyntenin and Cerebellin expression in the thalamus at E18.5**. *In situ* hybridization data from sagittal sections of the thalamus at E18.5 obtained from the devABA revealed differential patterns of expression of Calsyntenin (Clstn) 1 and 2, and Cerebellin (Cbln) 2 and 4. See text for details. Two representative and equivalent sections are shown for each gene, one lateral and the other medial (see Figure S7 for details on how the sections were selected).

***Netrin-Gs*** are GPI-anchored proteins related to netrins. They act as synaptic cell adhesion molecules via interactions with the transmembrane leucine-rich repeat proteins Netrin-G-ligand (NGL) -1 and -2 [93]. The devABA data match previous reports of broad Ntng1 expression across the dorsal thalamus and Ntgn2 expression restricted to the habenula (Figure 18) [94]. Ntng1 expression is not uniform, however; it displays significantly higher expression in some thalamic nuclei than others.

**Figure 18.**
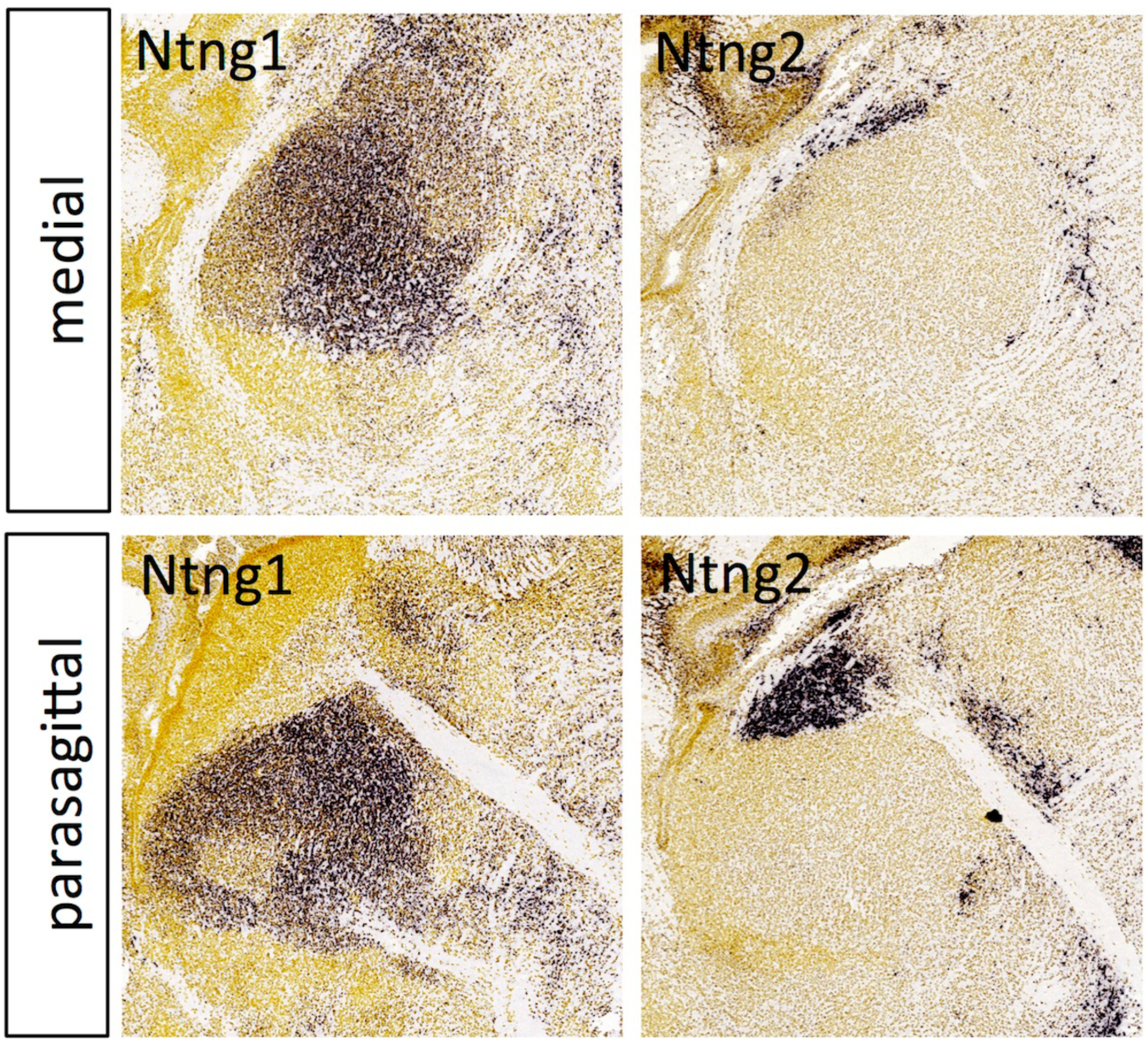
**Netrin-G expression in the thalamus at E18.5**. *In situ* hybridization data from sagittal sections of the thalamus at E18.5 obtained from the devABA revealed differential patterns of expression of Netrin-G (Ntng) 1 and 2. See text for details. Two representative and equivalent sections are shown for each netrin-G, one medial and the other adjacent medially (see Figure S7 for details on how the sections were selected).

***Miscellaneous genes:*** In addition to the gene families described above, there is a large set of miscellaneous genes encoding surface or secreted proteins with differential expression patterns (Figure 19). These include: Astn2, Cntnap4, Dlk1, Dner, Fzd7, Gpc3, Lrp8, Lypd1 (also known as Lynx2), Odz3 and Trp53i11 (also known as Tp53i11 or PIG11). These display varying degrees of specificity. For example, Fzd7 is highly restricted to what may become a single thalamic nucleus, while Gpc3, is expressed in a small number of scattered cells throughout the thalamus (and strongly in the reticular nucleus, not shown). Astn2, Cntnap4, Dlk1, Lypd1 all show quite selective expression in some developing nuclei and not others, while Dner, Lrp8 and Trp53i11 are expressed more widely, but still differentially. The Odz3 expression pattern documented in the devABA mirrored that seen in our own *in situ* hybridization experiments extremely faithfully (Figure S9).

**Figure 19.**
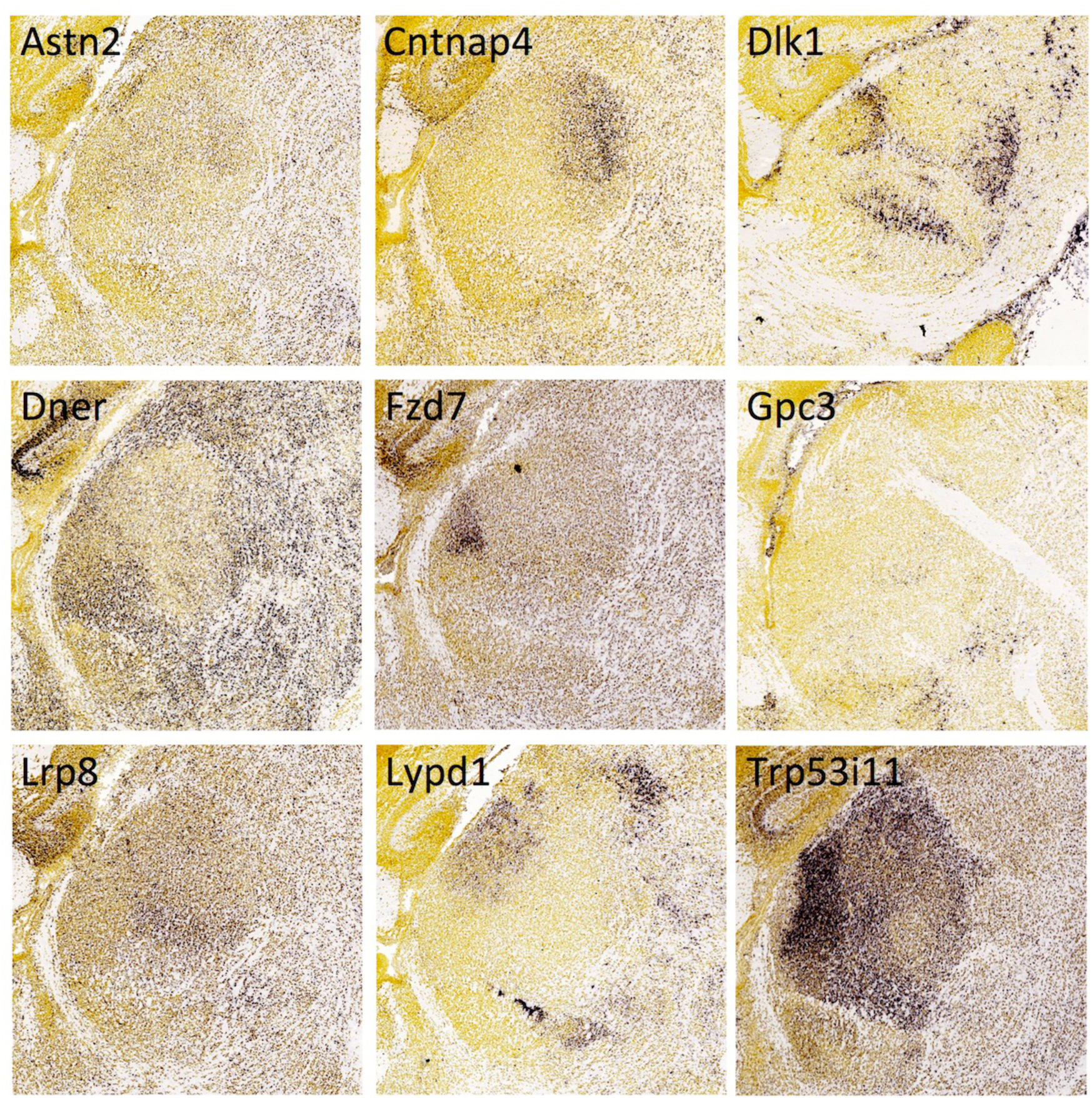
**Expression of genes encoding miscellaneous surface or secreted molecules in the thalamus at E18.5**. *In situ* hybridization data from sagittal sections of the thalamus at E18.5 obtained from the devABA revealed differential patterns of expression of Astrotactin2 (Astn2), Cntnap4, Delta-like-1 homolog (Dlk1), Delta/Notch-Like EGF Repeat Containing (Dner), Frizzled7 (Fzd7), Glypican-3 (Gpc3), Low-density lipoprotein receptor 8 (Lrp8), LY6/PLAUR Domain Containing 6 (Lypd1) and Trp53i11. See text for details. A representative and equivalent section is shown for each molecule (see Figure S7 for details on how the sections were selected). Dlk1 expression is shown laterally, while the other gene expression patterns are shown medially with Gpc3 more medial than the rest (parasagittal).

Lrp8 (low-density lipoprotein receptor 8) has been previously directly implicated in specifying thalamic connectivity. Mutation of *Lrp8*, along with the related gene *Vldlr (very low-density lipoprotein receptor)*, alters retinogeniculate innervation, through mechanisms that appear independent of the function of these proteins as Reelin receptors [11]. Mutation of *Cntnap4* affects synaptic outputs from GABAergic interneurons and ventral tegmental area dopaminergic neurons [95]. Gpc3 (Glypican-3) has been shown to bind to the synaptogenic molecule LRRTM4 [96] and, by analogy, with its close relative Gpc4, may actively specify synapse formation. Fzd7 (Frizzled-7) has not been directly implicated in synaptic connectivity but many other members of the Frizzled family have been [97], including Fzd5, which is required for the synapse-organising activity of Wnt7a [98]. Fzd5 is extremely selectively expressed in the parafascicular nucleus of thalamus [99], which we also find here (data not shown).

A variety of other functions in neural development or function have been shown for Astn2 (Astrotactin-2) [100], Dlk1 (Delta-like-1 homolog) [101], Dner (Delta/Notch-Like EGF Repeat Containing) [102] and Lypd1/Lynx2 (LY6/PLAUR Domain Containing 1) [103] but these genes have not been directly implicated in axon guidance or synaptic connectivity. The functions of Trp53i11 remain largely unknown.

***Growth factors and receptors.*** A variety of growth factors and receptors show selective or differential expression across the late embryonic thalamus (Figure 20). These include: Bmp3 (Bone morphogenetic protein-3), Igfbp5 (Insulin-like growth factor binding protein 5), Inhba (Inhibin/Activin, beta A subunit), Gfra1 and Gfra2 (GDNF Family Receptor Alpha 1 and 2), Nrn (Neuritin or Neuritin-1/Nrn1), Tgfb2 (Transforming growth factor, beta 2), Vgf (VGF nerve growth factor inducible) and Wif1 (WNT inhibitory factor 1).

**Figure 20.**
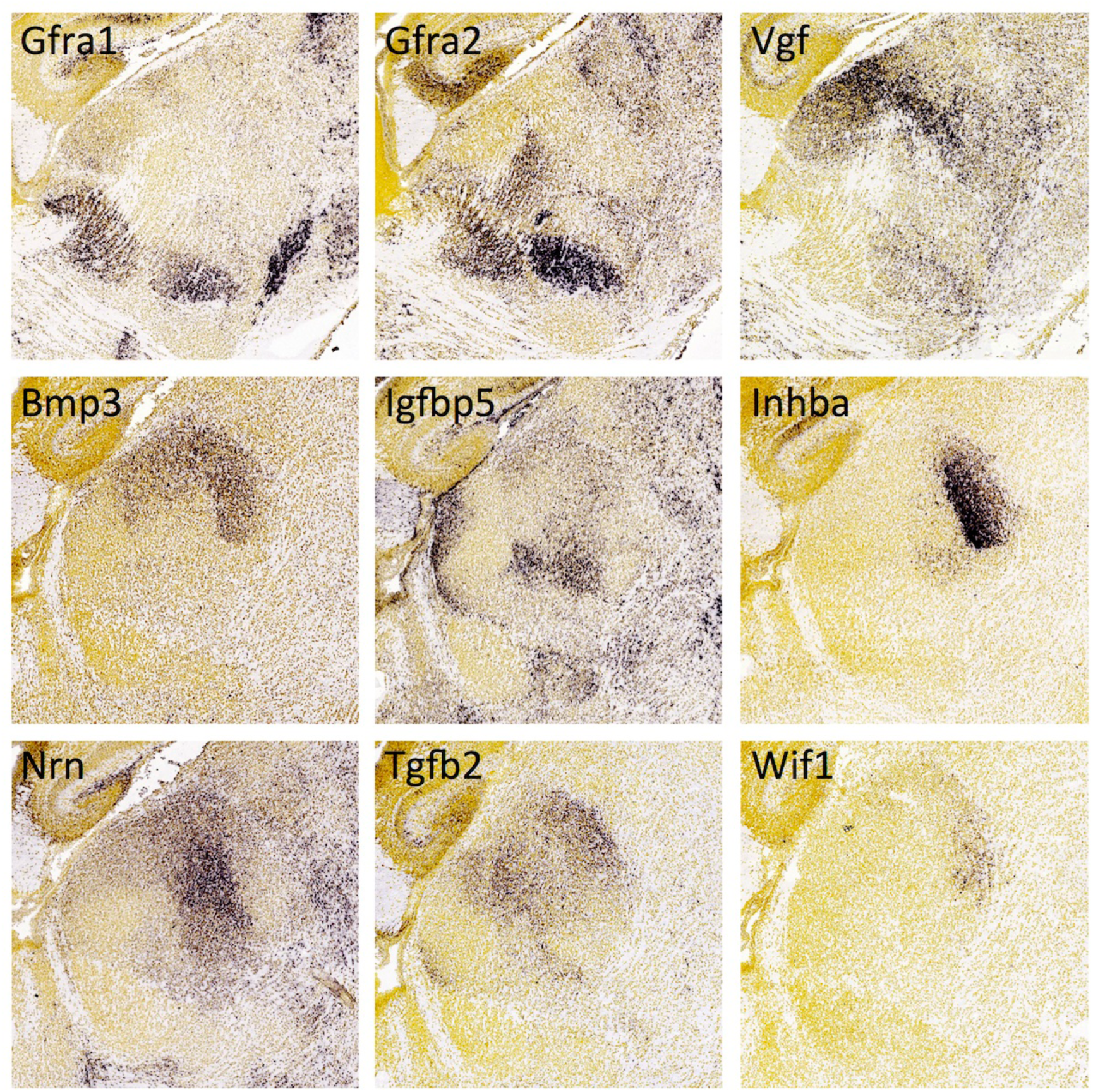
**Expression of growth factors and receptors in the thalamus at E18.5**. *In situ* hybridization data from sagittal sections of the thalamus at E18.5 obtained from the devABA revealed differential patterns of expression of Gfra1 and Gfra2 (GDNF Family Receptor Alpha 1 and 2), Vgf (VGF nerve growth factor inducible), Bmp3 (Bone morphogenetic protein-3), Igfbp5 (Insulin-like growth factor binding protein 5), Inhba (Inhibin/Activin, beta A subunit), Nrn (Neuritin or Neuritin-1/Nrn1), Tgfb2 (Transforming growth factor, beta 2) and Wif1 (WNT inhibitory factor 1). See text for details. A representative section is shown for each gene (see Figure S7 for details on how the sections were selected). Gfra1, Gfra2 and Vgf expression is shown laterally, while the other gene expression patterns are shown medially.

Differential expression of the GDNF receptors Gfra1, Gfra2 and Ret has been reported previously in postnatal rat thalamus, with notable overlap in the reticular nucleus [104], also seen in embryonic mouse [105]. These proteins have well defined roles in enteric nervous system formation but their functions in developing thalamus have not been elucidated. Selective expression of Neuritin-1 and VGF across thalamic nuclei has also been noted previously and these secreted molecules may play a role in regulating cortical differentiation in areas innervated by these thalamic axons [106]. Igfbp5, Inhba, Tgfb2 and Wif1 all have known roles in differentiation, proliferation and apoptosis in the developing brain but have not been directly implicated in axon guidance or synaptic connectivity. There are no described roles for Bmp3 in nervous system development.

## Discussion

Our dual screening approach has identified 82 genes encoding candidate connectivity labels in the developing thalamus. Several lines of evidence make these molecules highly plausible candidates to be involved in specifying some aspects of thalamic connectivity: (i) they encode proteins with motifs that are common in known connectivity molecules; (ii) some of them are members of gene families with already known functions in axon guidance or synaptic connectivity; (iii) they cluster with known axon guidance or synaptogenesis molecules based on their temporal pattern of expression in the embryonic thalamus; and (iv) they are selectively or differentially expressed across dorsal thalamic nuclei at stages when connectivity is being established.

The genes we have identified include multiple members of large families where some genes have been previously implicated, such as Ephrins and Eph receptors, cadherins and protocadherins, semaphorins and plexins, and Odz/teneurin genes, where we have expanded the number of candidates potentially involved. Our findings also highlight the selective expression in thalamus of a number of genes or gene families recently implicated in synaptic connectivity in other regions, including several Ig and LRR family members, calsyntenins, cerebellins and Netrin-Gs. In addition, we have found strikingly selective expression of many genes from diverse families not previously implicated in either thalamocortical development or neuronal connectivity.

The systematic nature of the devABA data provides the opportunity to collate and compare these expression patterns across gene families, and to draw some general inferences about the molecular logic specifying thalamic connectivity. The most striking aspect of these patterns is how unique they all are. Across these 82 genes, even when considering only one or two anatomical sections, we did not observe any two patterns that we would say are identical. Moreover, most of the differentiation of expression levels (at E15.5 or E18.5) reflects discrete nuclear divisions, rather than broad gradients across the whole dorsal thalamus, which, a priori, was just as likely an outcome. This indicates that thalamic nuclei are highly differentiated from each other, with each one displaying a unique repertoire of these molecules. There is, in addition, clear evidence of non-uniform gene expression within nuclei, reflecting the known sub-differentiation of these fields. Thus, while initial guidance of thalamic axons may be defined by general topography [reviewed in 9], later connectivity decisions are likely regulated by combinations of more spatially discrete labels.

There is indeed strong evidence for combinatorial interactions, both within and between the gene families identified. For example, co-expression of Ephrins and Eph-receptors in cis can alter responses to these proteins in trans [107]. Similar modulatory interactions in cis have been observed for Class 6 Semaphorins and Plexin-A proteins, thus generating combinatorial functional diversity [108-110]. Cadherins and protocadherins can also form heteromeric complexes, either within or across these two subfamilies [111,112], which can alter function, as with a cis interaction between Pcdh19 and Cdh2, which generates a novel trans-adhesive complex [113]. Ig superfamily members of the L1 and Cntn families can mediate homophilic adhesion, but also modulate signaling of semaphorin and Ephrin pathways in cis [71,114], while Cntns and Cntnaps also form functional complexes that regulate connectivity [reviewed in 115,116]. Multiple proteins in our dataset also interact with Neuroligin-Neurexin complexes implicated in synaptic development, including cerebellins [90], calsyntenins [39], Igsf9b [25], Mdga1 [78,79] and Gpc3 [96]. This pathway may thus represent a convergence point for combinatorial functions of multiple regulators of synaptogenesis, also including diverse members of the LRR superfamily and LAR protein tyrosine phosphatases not surveyed here [reviewed in 115,116].

Whether the genes we have identified actually encode connectivity labels will, of course, require functional experiments, but the list here is at least likely highly enriched for such molecules. It is by no means comprehensive, however, as both of our strategies had important limitations. The first approach concentrated on genes that are conserved from vertebrates to invertebrates, based on the simple rationale that many of the major families of axon guidance molecules are conserved. Such molecules may be expected to play more important and thus more obvious roles in nervous system development than newly evolved ones, with concomitantly stronger phenotypes when mutated. In addition, functional analyses of such gene families may be further simplified in invertebrate model systems, which often only have one gene copy, whereas mammals often have multiple genes with possibly redundant functions. Thus, while this strategy identified plausible genes that are highly amenable to functional analyses, it excluded the large fraction of the mammalian proteome that is not obviously conserved in invertebrates.

Our second strategy took advantage of the Allen Brain Atlas database of expression in the developing mouse brain [117]. This database enabled a systematic and quantitative comparison of expression profiles across seven embryonic and postnatal stages, which highlighted several clusters enriched for known or putative connectivity labels. We integrated annotations on protein topology, structural motifs and localization to better characterize the genes in this dataset and enable us to recognise possible surface labels. However, the dataset is limited to 2002 genes, which were selected on the basis of functional relevance to brain development and neurodevelopmental disorders, and which fall into several categories: (i) transcription factors, (ii) known regional or cell-type-specific markers, (iii) neurotransmitters and receptors, (iv) genes associated with patterning or axon guidance signaling pathways and (v) highly studied common drug targets, including GPCRs, ion channels, cell adhesion molecules and genes associated with neurodevelopmental disorders. While this set is thus itself likely enriched for connectivity labels, it is also certain that many others are not represented. For example, extracellular leucine-rich repeat (LRR) proteins are a major class of neuronal connectivity labels [118], with 135 such genes in the mouse genome [23], but only a small number are included in the devABA dataset.

In summary, the extensive set of genes identified here as candidate connectivity labels provides a strong starting point and context for functional analyses of single molecules and of their potential combinatorial interactions, which may coordinately specify the complex connectivity patterns of thalamic nuclei.

## Materials and Methods

### Mouse *in situ* hybridization

Digoxigenin (DIG)-labelled antisense cRNA probes for *in situ* hybridization were designed to encompass a section of coding sequence and 3’UTR >500bp in length. Briefly this involved TA cloning of PCR products into the TOPO vector (Invitrogen) and simultaneous synthesis and DIG-labelling of RNA transcripts from linearised vector using T7- or Sp6- RNA polymerase. Some cRNA probes were transcribed directly from the PCR products amplified with a sense primer and an antisense primer with T7 promoter sequence at its 5’-end. Primers used were:

Kirrel1_for TGACTATGGCACAGAGCCTA

Kirrel1_rev GAAGGAGAGGAAAGGGCA

Kirrel2_for CAAAAGAACTTGGTCCGGAT

Kirrel2_rev CACTTGGAGAGATGGATTCAC

Kirrel3_for TACGTGCAGTTTGACAAGGC

Kirrel3_rev AGGCTGCAAGGAATACAGAC

Igsf9_for AGACTCACCTCCTGCAAATC

Igsf9_T7Rev GCGGTAATACGACTCACTATAGGGCATTCCTGTTCAGCTCCCA

Igsf9b_for TGACACATTCCACAACGG

Igsf9b_T7Rev GCGGTAATACGACTCACTATAGGGCCCTTATTCCACTTCACCACAG

Sdk1_for CATCATACACTGTGGACCTG

Sdk1_T7rev GCGGTAATACGACTCACTATAGGGAGAAGTTCTGCACCCGTC

Sdk2_for AATGGTGGTTCTTAGTGGTCA

Sdk2_T7rev GCGGTAATACGACTCACTATAGGGCAAACGAAGGATGAGAACC

Odz1_for CATTCAGTGTGAGCTCCAGA

Odz1_T7rev GCGGTAATACGACTCACTATAGGGCTGACGCAAAGGCAGAGAT

Odz1_2_for TGAGGTCCAATATGAGATCC

Odz1_2_T7rev GCGGTAATACGACTCACTATAGGGCAACATCATAATCTCTTTGCCC

Odz2_for AGAGTCAAGCAAGCGAGAA

Odz2_T7rev GCGGTAATACGACTCACTATAGGGCGAGGTGGCAGCAGGTTAT

Odz2_2_for GTGCAATATGAGATGTTCCG

Odz2_2_T7rev GCGGTAATACGACTCACTATAGGGCAGTAAAGTGGACGAGCTTGG

Odz3_for GAAGAGTCAACAGTGGGAAGA

Odz3_T7rev GCGGTAATACGACTCACTATAGGGCTGCTACAGGAGAATCTGCAC

Odz3_2_for CAACCAAATCATTTCCACG

Odz3_2_T7rev GCGGTAATACGACTCACTATAGGGCTGGATTAGTTTGGTGAGCG

Odz4_for TTCAGAAGCAACTCAAGGC

Odz4_T7rev GCGGTAATACGACTCACTATAGGGCCAACACAGTCAGGAATACGG

Odz4_2_for AGAAGGAGCTGAAGGTGG

Odz4_2_T7rev GCGGTAATACGACTCACTATAGGGCCACGATGCTGTTGCTACTC

Neto1_for GGCTCCAGAACTGTGTATATCC

Neto1_T7rev GCGGTAATACGACTCACTATAGGGCGAGTCTTGCCGAAGGAATA

Neto1_2_for AGAAGTCAGTGCAGTGTGG

Neto1_2_T7rev GCGGTAATACGACTCACTATAGGGCGAGCTGCTTCCGTCATAG

Neto2_for CATCAGGAATTGTCTTGGTC

Neto2_T7rev GCGGTAATACGACTCACTATAGGGCCTGCTCTGCCTGACTTAACA

Neto2_2_for GAATCAAGCACATTCCTGC

Neto2_2_T7rev GCGGTAATACGACTCACTATAGGGCGGATCACTCCGACTCCTG

Clstn1_for CACCCTGAACATCGATCC

Clstn1_T7rev GCGGTAATACGACTCACTATAGGGCTCACCCTCTGTCGACAAG

Clstn2_for GCCACTGTCGTCATCATTATC

Clstn2_T7rev GCGGTAATACGACTCACTATAGGGCATGGTGACATCATCTCCAGA

Clstn3_for TCCCAGTCATGTGCTCAGT

Clstn3_T7rev GCGGTAATACGACTCACTATAGGGCAGTAGAGAAGGACACAGCAGC

*In situ* hybridization was carried out on vibratome-sectioned C57BL6 mouse brains (Jackson Laboratories) as described in [119] with modifications. To obtain brains at E15, timed pregnancy mice were sacrificed by cervical dislocation and embryos were fixed and stored in 4% paraformaldehyde (PFA)-PBS at 4°C until use. For P0, brains were dissected out prior to immersion in 4% PFA-PBS -at 4°C. P9 brains were fixed by perfusion followed by immersion in 4% PFA-PBS at 4°C. Fixed brains were embedded in 3% agarose and 70-μm sections were obtained on a vibratome (VT1000S Leica). Sections were washed twice in PBST (PBS containing 0.1% Tween-20), permeabilised in RIPA buffer (150mM NaCl, 50mM Tris-HCl pH 8.0, 1mM EDTA, 1% Nonidet-P40, 0.5% sodium deoxycholate, 0.1% SDS) and postfixed in PBS containing 4% PFA and 0.2% gluteraldehyde. Hybridisation was performed in a humidified environment overnight at 65°C with 1μg/ml labeled probe in hybridisation buffer (50% formamide, 5X SSC pH4.5, 1% SDS, 50μg/ml yeast tRNA, 50μg/ml heparin). Posthybridisation washes were completed at 65°C using solution I (50% formamide, 5X SSC pH4.5, 1% SDS) and solution III (50% formamide, 2X SSC pH4.5, 0.1% Tween-20) and at room temperature using TBST (TBS containing 1% Tween-20). Brain sections were incubated for >1hr in blocking buffer (TBST, 10% heat-inactivated sheep serum). Immunodetection was carried out in blocking buffer at 4°C overnight using an alkaline phosphatase-conjugated anti-digoxigenin antibody (Roche) at a 1:2000 dilution. Following antibody incubation extensive TBST washes were performed. Sections were equilibrated in NTMT (100mM Tris-HCl pH 9.5, 100mM NaCl, 50mM MgCl_2_, 1% Tween-20) prior to colourimetric detection using 2μl/ml NBT/BCIP (Roche) in NTMT. Sections were mounted on Superfrost glass slides (VWR international) and analysed with an Olympus IX51 microscope.

### Database pipeline

The database pipeline was used as described in [23]. In summary, the Ensembl database, the International Protein Index and the Mammalian Gene Collection were used to build proteomes for human, mouse, worm and fly. Through automated and manual curation duplicates were removed and only the longest isoforms were kept. This resulted in non-redundant data sets for mouse, human, worm, and fly containing 85991, 74866, 22698, and 16857 sequences, respectively. From an all-against-all Blast search the top 200 hits were used as input for Markov clustering with the MCL program. The output was combined with rich annotation, including predictions of motifs and GPI links, as well as transmembrane and signalling sequences.

### Data acquisition and preprocessing

The Allen Brain Atlas (ABA) provides a survey of gene expression in the brain from mid-gestation through aging in different species. Of interest to us is the data on the mouse brain. The detailed process of mouse brain data acquisition is explained in Lein et al. [120]. Briefly, *in situ* hybridization (ISH) data was recorded from coronal or sagittal sections and quantified per unit from a given spatial location for each gene (expression values per voxels). Voxels were assigned regional labels through age-matched anatomic reference atlases. The developmental dataset on the ABA portal (www.brain-map.org) comprises expression values of 2002 genes obtained from ISH images in the sagittal plane across four embryonic (E11.5, E13.5, E15.5 and E18.5) and three early postnatal ages (P4, P14 and P28) [117]. These time-points are developmentally relevant to regional specification/patterning (E11.5), axon pathfinding (E13.5, E15.5 and E18.5), synaptogenesis (P4 and P14) and cortical plasticity (P28). One animal was used per time-point per gene. Per-region data was quantified from ISH images by combining all voxels with the same regional label. Liscovitch and Chechik [121] extracted data from 36 anatomically delineated regions of the developing brain, which encompass the entire brain. The data is readily available for download at http://chechiklab.biu.ac.il/~lior/cerebellum.html. The data is expressed as expression density. For each brain region R, the expression density is defined as the sum of expressing pixels in R divided by the total number of pixels that intersect R (taken from the Technical White Paper: Informatics Data Processing for the Allen Developing Brain Atlas found at http://developingmouse.brain-map.org/docs/InformaticsDataProcessing.pdf). Each set of values is dependent on specific gene expression. We normalized the data to reflect variations of expression for each gene. A value of 1 indicates average expression levels whereas a value below or above 1 indicates decreased or increased expression levels compared to average, respectively. With this normalised data, we generated the developmental matrix of 1996 genes x 7 time-points as 6 genes had no expression values throughout. These genes are Chrna1, Hoxd9, Lef1, Nrp2, Pbx1 and Rnd2.

The genes of the developmental dataset fall into one of these five categories (taken from the Technical White Paper: Allen Developing Mouse Brain Atlas found at http://developingmouse.brain-map.org/docs/Overview.pdf):

1. *Transcription factors*. Approximately one third of the genes are transcription factors, with extensive coverage of homeobox, basic helix-loop-helix, forkhead, nuclear receptor, high mobility group and POU domain genes.
2. *Neuropeptides, neurotransmitters, and their receptors*. Extensive coverage of genes in dopaminergic, serotonergic, glutamatergic, and gabaergic signaling, as well as neuropeptides and their receptors.
3. *Neuroanatomical marker genes*. Characterizing region- or cell type specific marker genes over development can provide information about the origins of a brain region or cell type, and may help to identify precursor regions at earlier time points.
4. *Gene ontologies/signaling pathways relevant to brain development*. Gene ontologies include axon guidance, receptor tyrosine kinases and their ligands. Pathways include Wnt signaling and Notch signaling pathways.
5. *Genes of general interest*. This category includes highly studied genes such as common drug targets, ion channels, cell adhesion, genes involved in neurotransmission, G-protein-coupled receptors, or involved in neurodevelopmental diseases, which are expressed in the brain in the adult and/or in development.

We screened protein products for cellular localization with an emphasis on the extracellular compartment. More specifically, we looked for signal peptides, transmembrane domains and glycophosphatidylinositol (GPI)-anchors. Since genes were listed with their Allen gene name in the dataset, we looked for the MGI symbol and Ensembl gene identifiers (through http://www.ncbi.nlm.nih.gov/gene) to subsequently find their protein counterparts. Where multiple isoforms exist, the longest sequence was used for further analyses. We screened Fasta format sequences through PSORTII to find cellular localization of the protein products and the number of predicted helices of transmembrane proteins (http://psort.hgc.jp)[122,123]. PSORTII predicted protein localization in the following compartments: cytoplasm, cytoskeleton, endoplasmic reticulum, Golgi, mitochondria, nucleus, plasma membrane, peroxisomes, vacuoles and vesicles of the secretory system. In addition, we ran the SignalP prediction program to determine whether a protein contains a signal peptide or not - indicative of extracellular localization (http://www.cbs.dtu.dk/services/SignalP/, version 4.1)[124,125]. A GPI modification site prediction, the big-PI predictor was run to identify protein products with potential GPI-anchors (http://mendel.imp.ac.at/sat/gpi/gpi_server.html, version last modified June 17^th^ 2005) [126-129]. When any of the three programs suggested extracellular localization, the protein was deemed found in the extracellular compartment. The TMHMM program was run to assess the presence and number of transmembrane domains (http://www.cbs.dtu.dk/services/TMHMM/, version 2.0)[130,131]. When the number of transmembrane domains varied between PSORTII and TMHMM results, manual curation was performed (unless both programs picked up over two transmembrane domains and the protein was simply considered multi-pass). PFAM (http://pfam.xfam.org)[132] and SMART (http://smart.embl-heidelberg.de, version 2.12.1 released on August 6^th^ 2012)[133,134], motif recognition programs, allowed screening for important protein domains, which can be amalgamated through InterPro (http://www.ebi.ac.uk/interpro/search/sequence-search, version 49.0 released on November 20^th^ 2014)[135]. For identification of important protein domains we retrieved PFAM, SMART, Interpro and SignalP information through BioMart version 0.7 [136].

Finally, we updated the gene annotations of the dataset from Thompson et al. [117]. Thompson and colleagues had 1385 genes out of the 2002 falling into one or more of these categories: axon guidance pathway, cell adhesion, receptor tyrosine kinases (RTKs) and their ligands, Notch signaling, Wnt signaling, transcription factor activity, basic helix-loop-helix transcription factor, forkhead transcription factor, homeobox transcription factor, nuclear receptor, POU domain genes, neurotransmitter pathway, ion channel, and G-protein coupled receptors (GPCRs). We screened genes for the following gene ontology (GO) term annotations (http://www.geneontology.org) [137] through MartView:

1. Axon Guidance GO:0007411;
2. Synapse GO:0045202;
3. Patterning: - Regionalization GO:0003002;
  - Pattern Specification Process GO:0007389;
  - Neural Tube Patterning GO:0021532;
  - Rostrocaudal Neural Tube Patterning GO:0021903;
4. Transcription Factor Activity:
  - Protein Binding Transcription Factor Activity GO:0000988;
  - Nucleic Acid Binding Transcription Factor Activity GO:0001071;
  - Sequence-Specific DNA Binding Transcription Factor Activity GO:0003700.

Since many genes had more than one annotation, we grouped them as follows:

- Group 1: axon guidance pathway and cell adhesion;
- Group 2: synapse;
- Group 3: receptor tyrosine kinases and their ligands, and patterning;
- Group 4: neurotransmission pathway (GPCRs, ion channels, gap junctions);
- Group 5: chromatin and transcription factor activity
- Group 6: other (cytoskeleton, extracellular matrix, myelin, metabolic enzymes and signal transduction) and unannotated.

We only had 206 unannotated genes. Each gene belonged to only one group. When a gene fell into more than one group, we prioritised the groups as presented.

### Clustering analyses

We performed *k*-means clustering with the Euclidean distance method on the developmental dataset using the clustering module integrated in ArrayPipe (http://www.pathogenomics.ca/arraypipe - version 1.7)[138]. Each clustering analysis was iterated a thousand times.

### Enrichment analyses

We performed chi square statistics to determine whether our developmental dataset was normally distributed in regards to cellular localisation and groups across the 10 clusters obtained during *k*-means clustering analyses or enriched in particular clusters. Statistical significance was determined at p-value < 0.05. Where multiple tests were performed on the same dataset, the Bonferroni correction was taken into account.

## Acknowledgments

The authors would like to thank Dr. Jackie Dolan for help in managing animal colonies and all the animal facility staff at Trinity College Dublin. This work was supported by grants to KJM from the Wellcome Trust (075264/A/04/Z) and Science Foundation Ireland (07/IN.1/B969 and 09/IN.1/B2614) and to OBB from the Fonds de recherche Santé Québec (23980)

## Supplemental Tables

**Table S1. Conserved candidate connectivity labels.** Genes are organized in clusters defined by TRIBE-MCL, as described [23]. Genes from Drosophila melanogaster are in blue, from Caenorhabditis elegans in green and from mouse in black. Signal peptides were predicted using SignalP, transmembrane domains by a consensus between TMHMM and HMMTOP and protein motifs by SMART and Pfam, also as described [23].

**Table S2. Developmental dataset.** Absolute values of expression densities for 2002 genes from the devABA database per gene per time-point are shown.

**Table S3. Developmental dataset and *k*-means clustering.** Normalised expression densities per gene per time-point (columns B-H). Thalamic expression densities were obtained for 1996 genes. Heatmap’s 3 colour scale of gene expression data: 0.2, red; 1, white; blue, 5. Normalised gene expression was clustered by similar patterns of expression using *k*-means Euclidean distance 6-18, with 1000 iterations (columns I-U). Clusters are sorted at *k* = 10, followed by *k* = 11, followed by *k* = 12 and so on until *k* = 18. Three colour scale of cluster number: red < yellow < green, for better visualisation.

**Table S4. Developmental dataset protein characteisation**. Each gene was matched to its MGI symbol and Ensembl gene identifier. From these, protein sequences were obtained - where multiple isoforms were found, the longest sequence was used for further analyses. Fasta format sequences were screened through PSORT II for cellular localization (PSORT II prediction and probability: gol, Golgi; cyt, cytoplasmic; csk, cytoskeletal; end, endoplasmic reticulum; ext, extracellular, including cell wall; mit, mitochondrial; nuc, nuclear; per, peroxisomal; pla, plasma membrane; ves, vesicles of secretory system). PSORT II uses two-fold *k-nearest* neighbor (k-NN) algorithm with *k1* = 9 and *k2* = 23 [122]. Screens for number of predicted transmembrane domains (TMs) with PSORT II were also performed. SignalP detected signal peptides (sigp) on proteins. Big-PI predictor tested different potential GPI-modification sites and returned the GPI best score, GPI profile score, GPI profile independent score and GPI quality [126-129]. GPI quality of P or S indicated a potential GPI-modification site (P: predicted; S: second predicted site, more than one predicted site). TMHMM screened for the presence (Transmembrane Domain) and number of transmembrane domains (PredHel) [130,131]. Columns AE, AF and AG show SMART ID, PFAM ID and InterPro short description, respectively. Genes were screened for additional GO terms (see methods).

**Table S5. Developmental dataset protein localisation and functional annotations.** Normalised expression densities per gene per time-point (Columns B-H). Heatmap’s 3 colour scale of gene expression data: 0.2, red; 1, white; 5, blue. Data is sorted by cluster number at *k* = 10 followed by gene name (Column I). Each gene was matched to its MGI symbol (Column J). Protein localisation was assessed after combining the data from the different screens performed as either in the Golgi (gol), cytoplasmic (cyt), cytoskeletal (csk), in the endoplasmic reticulum (end), extracellular including cell wall (ext), mitochondrial (mit), nuclear (nuc), peroxisomal (per), on the plasma membrane (pla) or in vesicles of secretory system (ves; Column K). Moreover, when the protein was deemed in the extracellular compartment, it was found to be either GPI-achored (anchored), secreted, single-pass transmembrane (single-pass) or multi-pass transmembrane (multi-pass; Column L). Functional assessment was grouped in six different categories: 1) axon guidance pathway and cell adhesion; 2) synapse; 3) receptor tyrosine kinases and their ligands, and patterning; 4) neurotransmission pathway (GPCRs, ion channels, gap junctions); 5) chromatin and transcription factor activity; and 6) other (cytoskeleton, extracellular matrix, myelin, metabolic enzymes and signal transduction) and unannotated (Column M). Three colour scale of cluster number and group number: red < yellow < green, for better visualisation (Columns I and M). See text for details.

**Table S6. Known and potential connectivity genes.** Genes are listed by cluster and grouped into those with known roles in thalamocortical connectivity, known functions in axon guidance of synaptogenesis more generally or no such known functions (Other).

## Supplemental Figures

**Figure S1.**
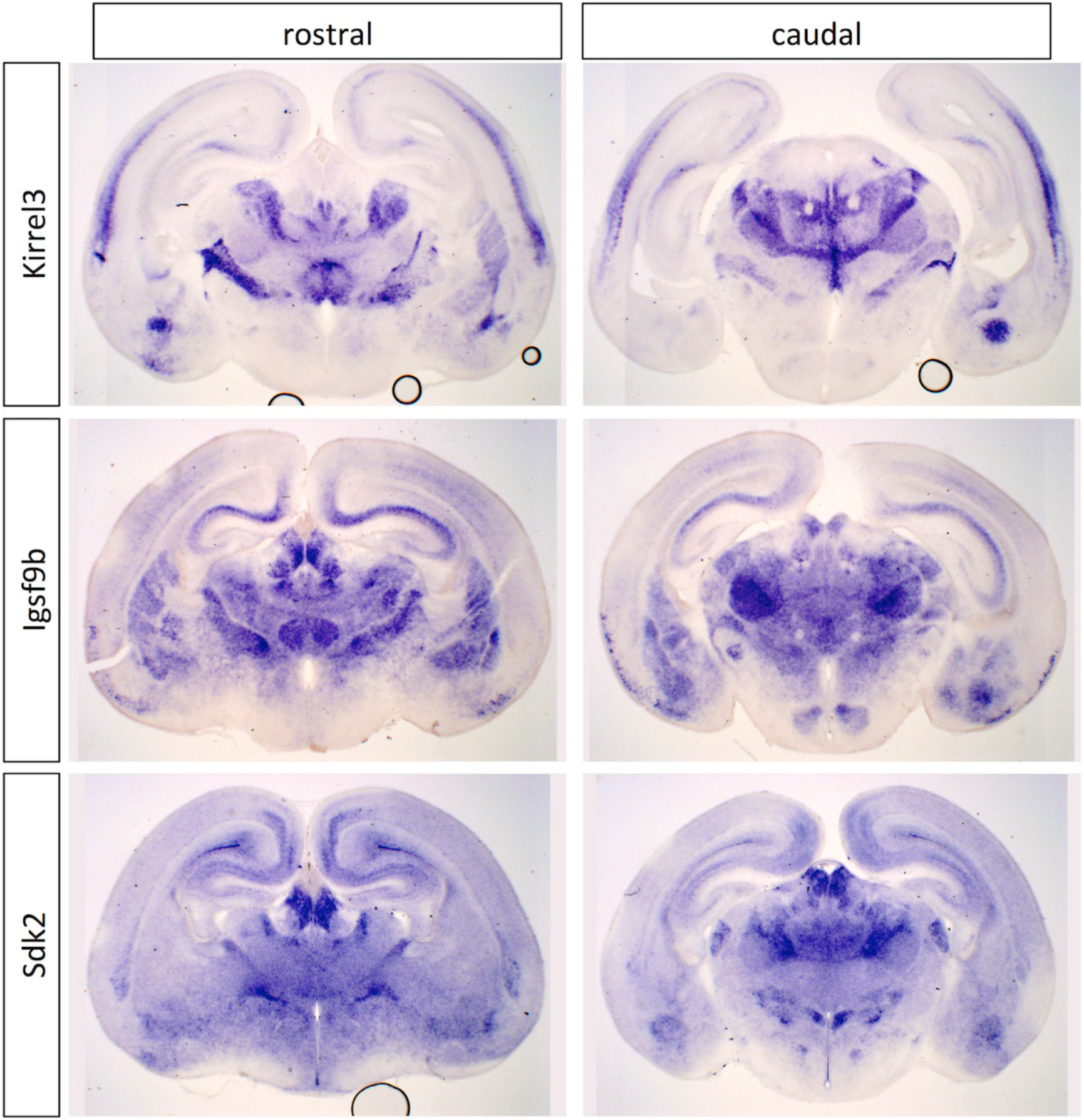
***In situ* hybridization patterns for immunoglobulin superfamily genes across entire brain at P0.** Two coronal sections are shown for Kirrel3, Igsf9b and Sdk2, one rostral and one more caudal.

**Figure S2.**
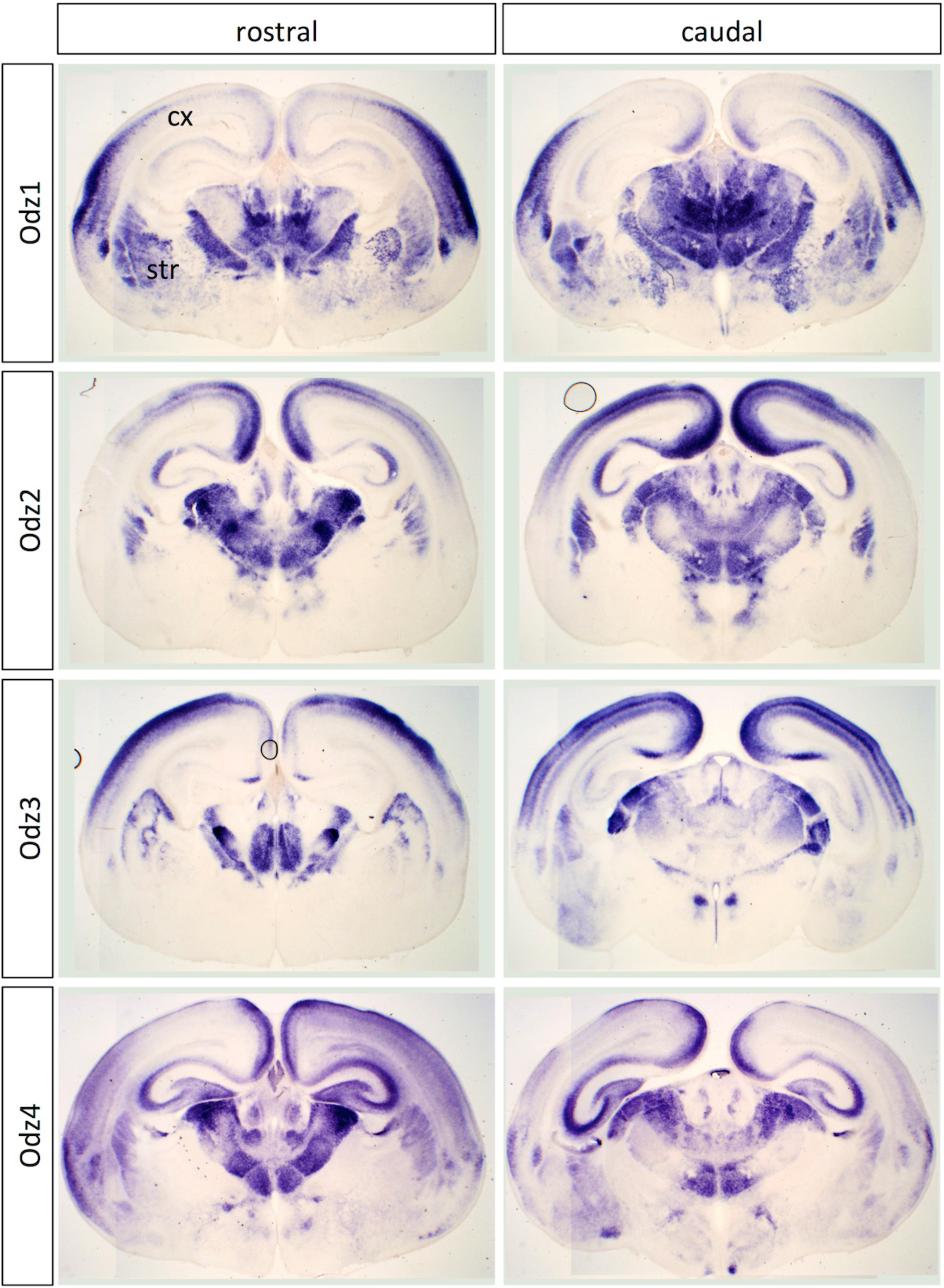
***In situ* hybridization patterns for Odz family genes across entire brain at P0.** Two coronal sections are shown for Odz1, Odz2, Odz3 and Odz4, one rostral and one more caudal. In addition to restricted expression in dorsal thalamus, there is also graded expression of Odz genes across cortex (cx) and striatum (str) in differing patterns.

**Figure S3.**
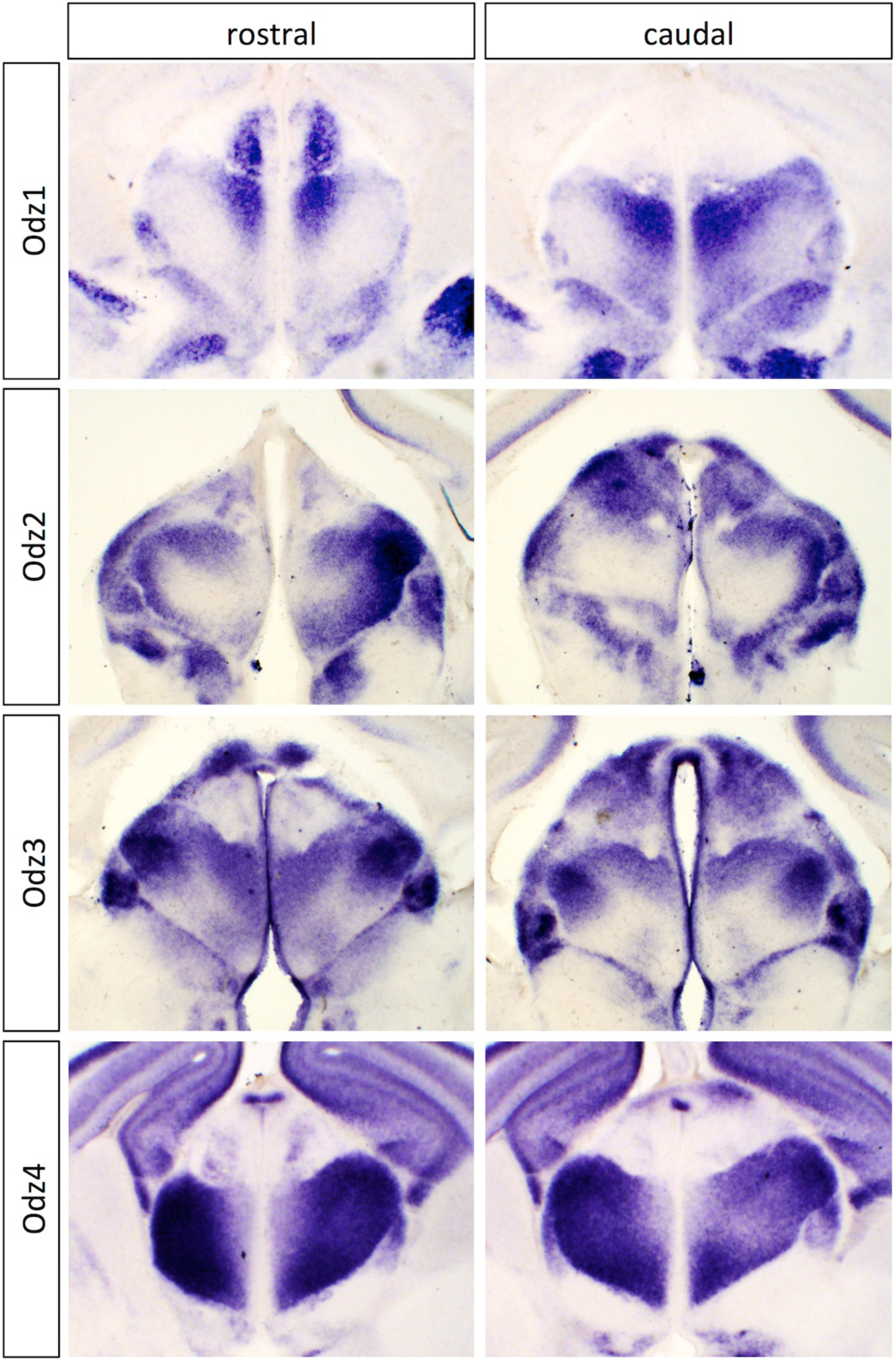
***In situ* hybridization patterns for Odz family genes in dorsal thalamus at E15.5.** Two coronal sections are shown for Odz1, Odz2, Odz3 and Odz4, one rostral and one more caudal. Differential expression across the dorsal thalamus is already evident at this stage. The corresponding entire brain sections are shown in Figure S4.

**Figure S4.**
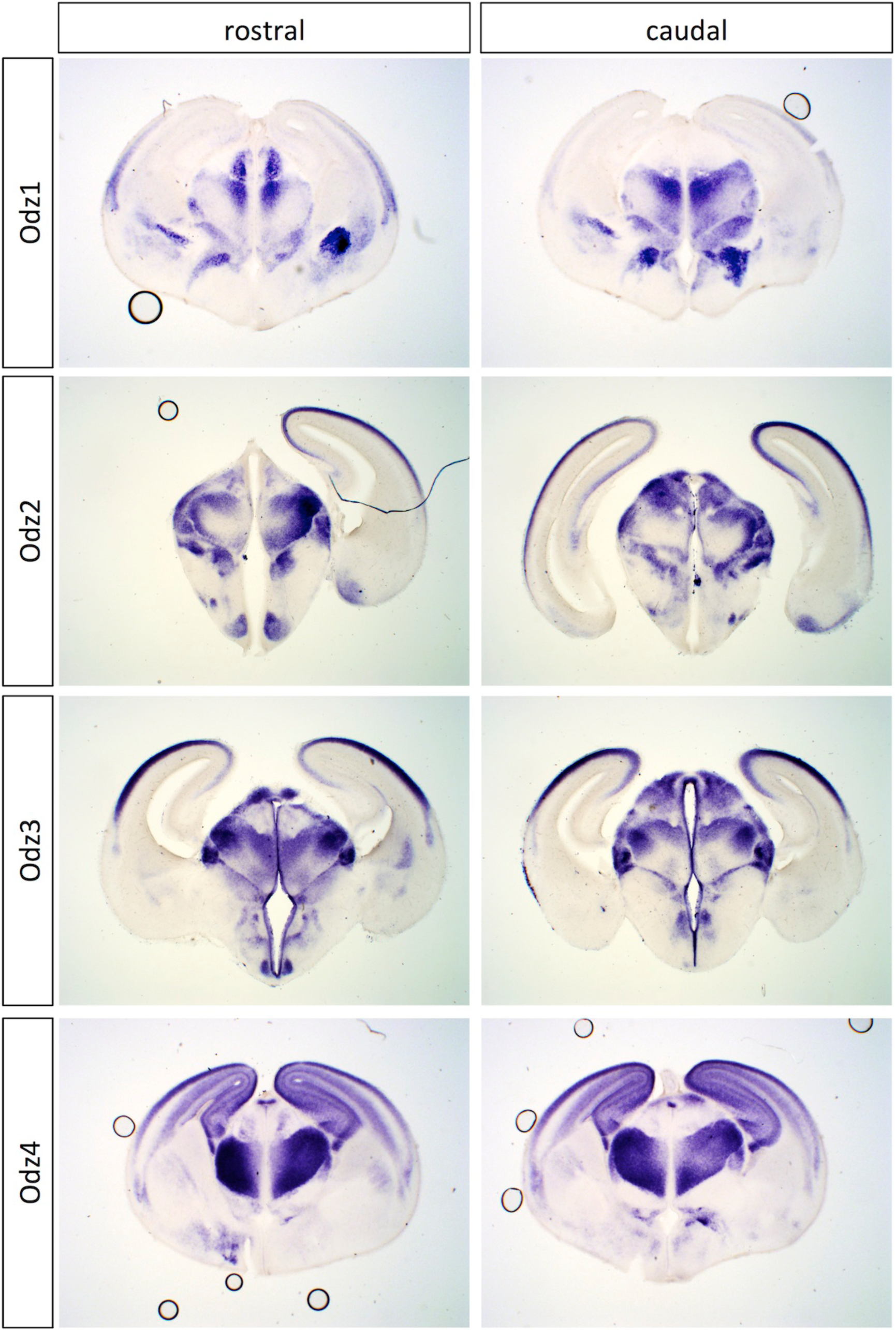
***In situ* hybridization patterns for Odz family genes across entire brain at E15.5.** Two coronal sections are shown for Odz1, Odz2, Odz3 and Odz4, one rostral and one more caudal.

**Figure S5.**
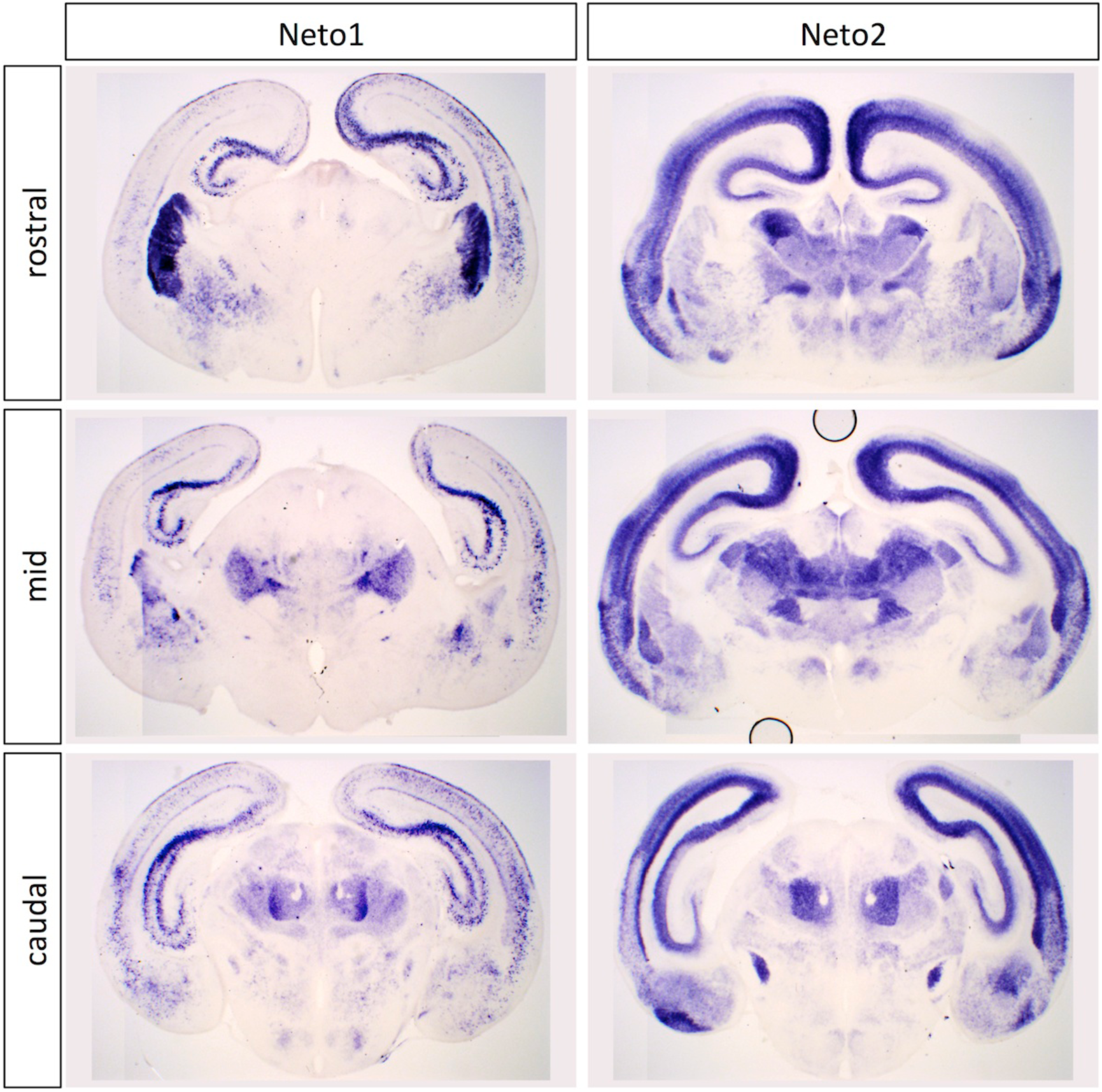
***In situ* hybridization patterns for Neto family genes across entire brain at P0.** Three coronal sections are shown for Neto1 and Neto2, one rostral, one at an intermediate level (mid) and one more caudal.

**Figure S6.**
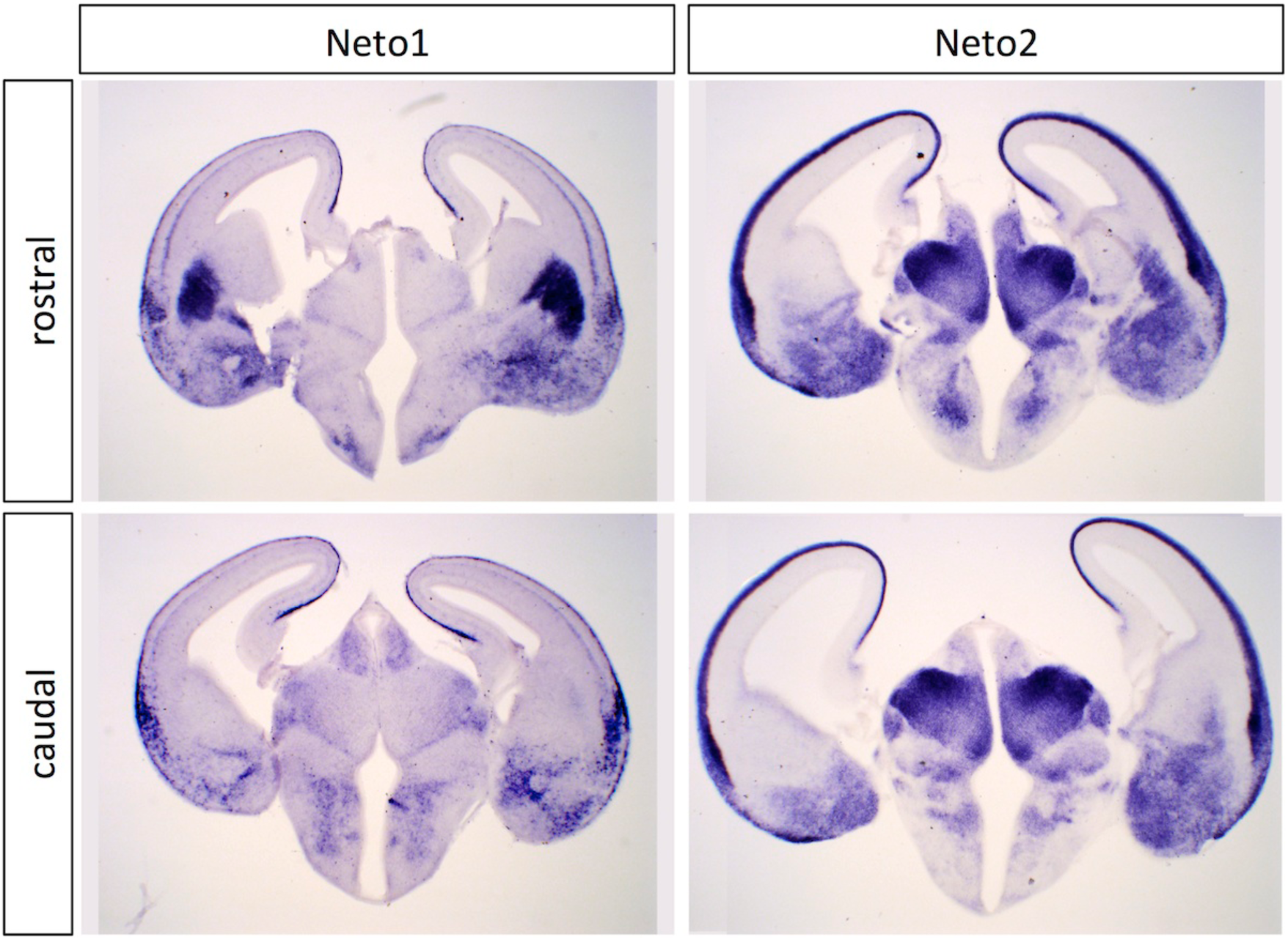
***In situ* hybridization patterns for Neto family genes across entire brain at E15.5.** Two coronal sections are shown for Neto1 and Neto2, one rostral and one more caudal. Differential expression across the dorsal thalamus is already evident at this stage.

**Figure S7.**
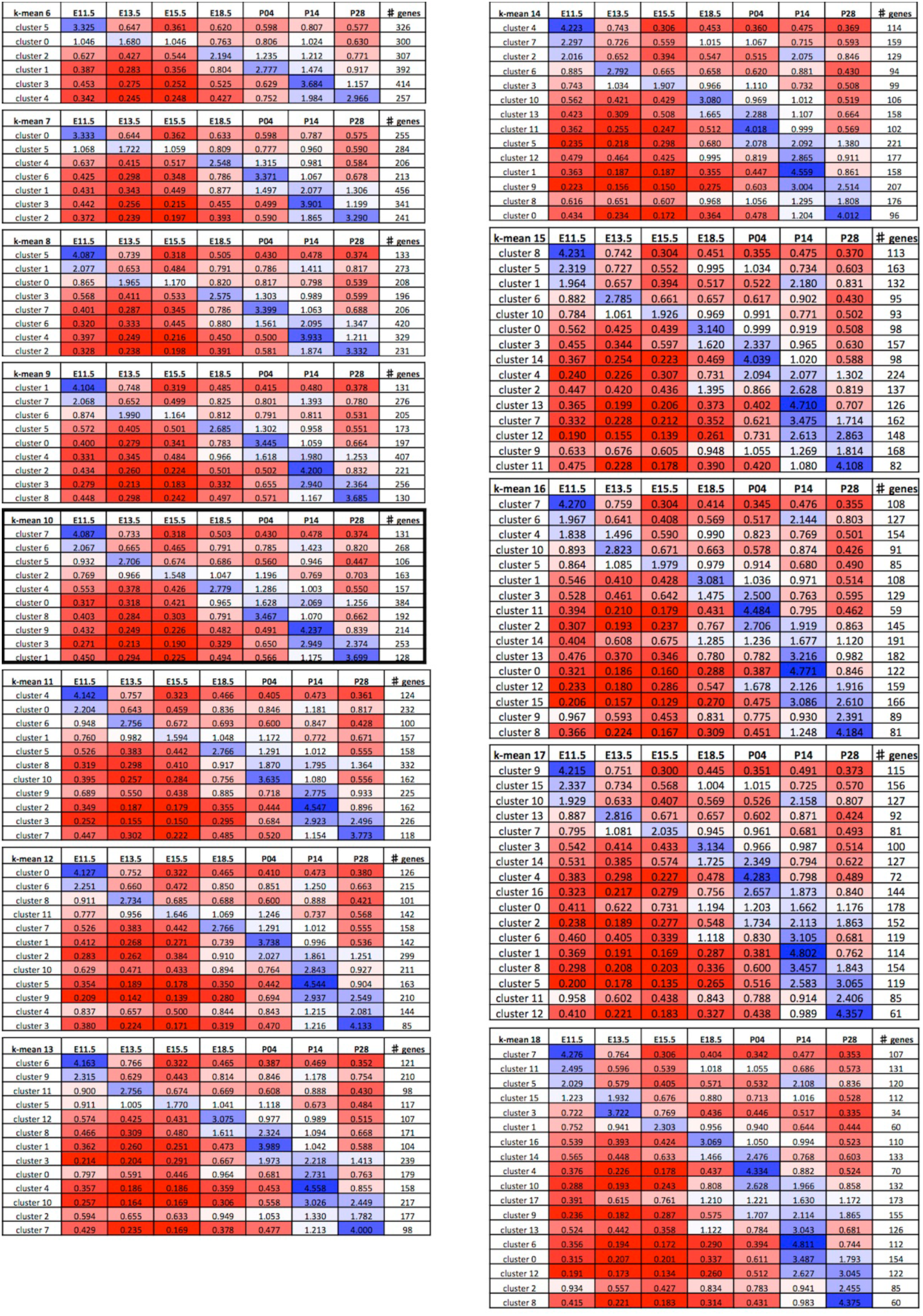
**Summaries of expression profiles across *k*-means values.** Normalised expression densities were averaged per cluster to see the trends of expression for the results of all clustering analyses from *k* = 6 to 18 (*k* indicated at top left corner of each table). Clusters were organised chronologically with early peaks of expression at the top and later peaks of expression at the bottom. Heatmap’s 3 colour scale of gene expression data: 0.2, red; 1, white; 5, blue. *k* = 10 was used for further analyses; the corresponding tableframe is in bold.

**Figure S8.**
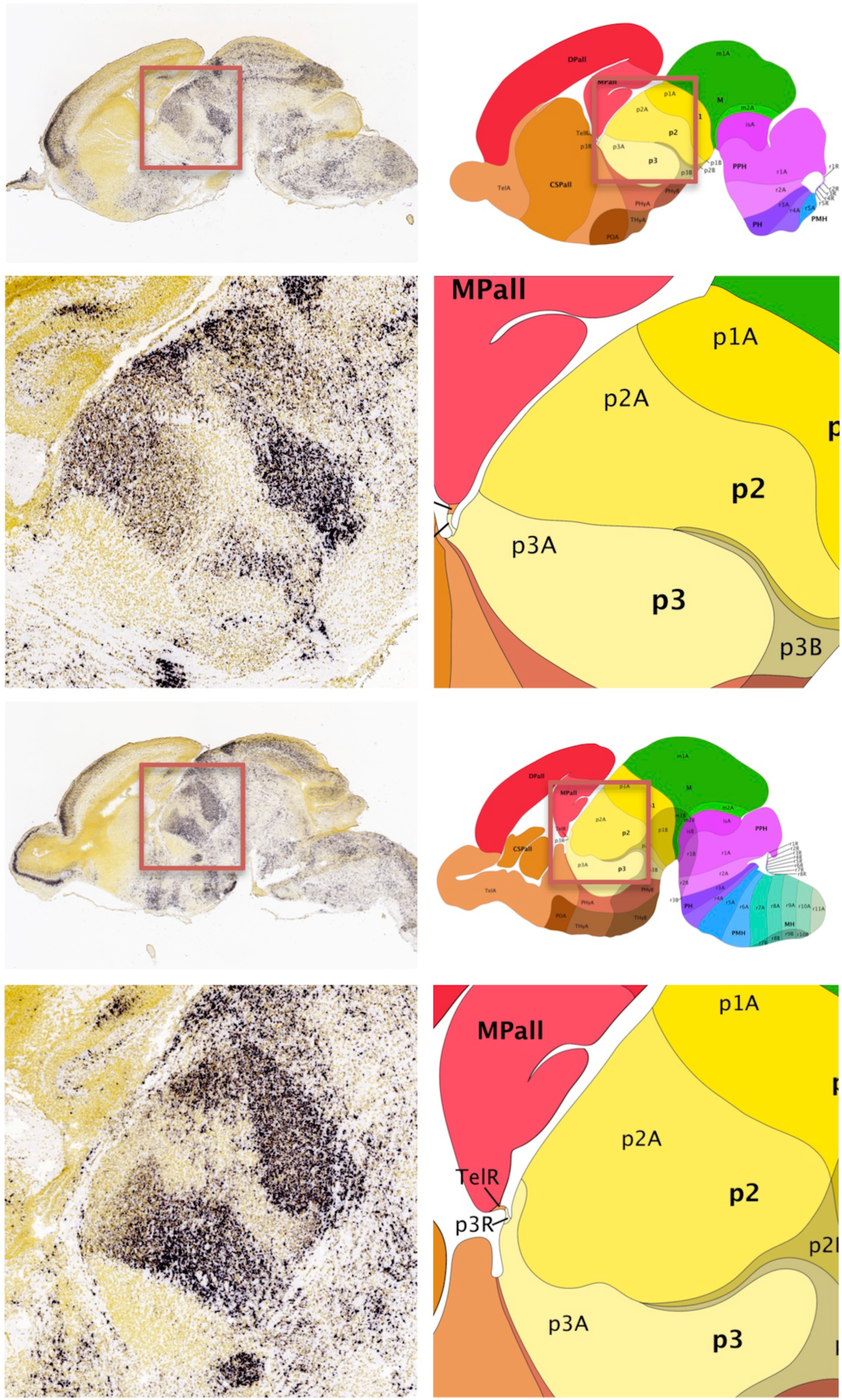
**Image extraction of thalamic gene expression from the devABA.** *In situ* hybridization data from a sagittal section of the thalamus at E18.5 obtained from the devABA (A). Corresponding section from the anatomic reference atlas (B). Higher magnifications of the thalamus from squared regions in A and B, respectively (C and D). p1, prosomere 1 (pretectum and pretectal tegmentum); p1A, alar plate of prosomere 1; p1B, basal plate of prosomere 1; p2, prosomere 2 (thalamus and thalamic tegmentum); p2A, alar plate of prosomere 2; p2B, basal plate of prosomere 2; p3, prosomere 3 (prethalamus and prethalamic tegmentum); p3A, alar plate of prosomere 3; p3B, basal plate of prosomere 3. Note that voxels were assigned regional labels for thalamus and thalamic tegmentum separately.

**Figure S9.**
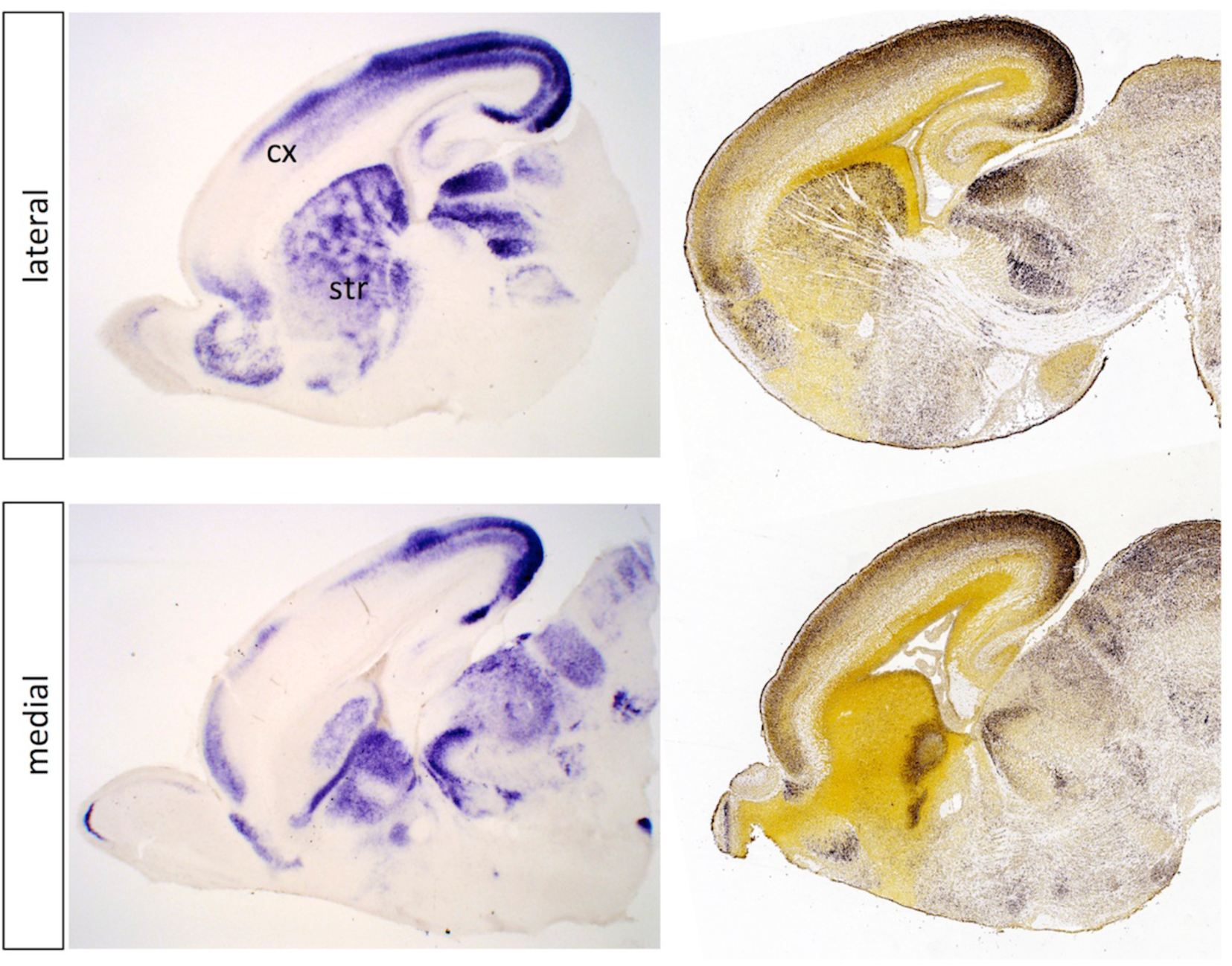
**Odz3 *in situ* hybridization compared to devABA data.** Two sagittal sections are shown, one lateral and one more medial. Our *in situ* hybridizations are on sections from P0 nenonates, while the devABA sections are from E18.5 embryos. Despite this difference of about a day, there is strikingly good correspondence in expression patterns across the dorsal thalamus, striatum (str), cortex (cx) and other brain regions.

